# Pleomorphic Adenoma Gene 1 Is Needed For Timely Zygotic Genome Activation and Early Embryo Development

**DOI:** 10.1101/305375

**Authors:** Elo Madissoon, Anastasios Damdimopoulos, Shintaro Katayama, Kaarel Krjutškov, Elisabet Einarsdottir, Katariina Mamia, Bert De Groef, Outi Hovatta, Juha Kere, Pauliina Damdimopoulou

## Abstract

Pleomorphic adenoma gene 1 (PLAG1) is a transcription factor involved in cancer and growth. We discovered a *de novo* DNA motif containing a PLAG1 binding site in the promoters of genes activated during zygotic genome activation (ZGA) in human embryos. This motif was located within an Alu element in a region that was conserved in the murine B1 element. We show that maternally provided *Plag1* is essential for timely mouse preimplantation embryo development. Heterozygous mouse embryos lacking maternal *Plag1* showed disrupted regulation of 1,089 genes, spent significantly longer time in the 2-cell stage, and started expressing *Plag1* ectopically from the paternal allele. The *de novo* PLAG1 motif was enriched in the promoters of the genes whose activation was delayed in the absence of *Plag1*. Further, these mouse genes showed a significant overlap with genes upregulated during human ZGA that also contain the motif. By gene ontology, the mouse and human ZGA genes with *de novo* PLAG1 motifs were involved in ribosome biogenesis and protein synthesis. Collectively, our data suggest that PLAG1 affects embryo development in mice and humans through a conserved DNA motif within Alu/B1 elements located in the promoters of a subset of ZGA genes.

## INTRODUCTION

Early preimplantation embryo development is dependent on zygotic genome activation (ZGA) (1, 2). Transcription from the newly formed zygotic genome starts gradually already in the one-cell embryo, and a major increase in transcriptional output, known as major ZGA, takes place during the 2-cell (2c) stage in mice and during the 4c-to-8c transition in humans (1, 2). Both minor and major ZGA are essential for cleavage stage development in the mouse (3, 4) and thus needed for the formation of a blastocyst capable of implanting to the uterine endometrium. Therefore, knowledge about the gene expression program during the first stages of embryonic development and regulation of ZGA will help discover factors controlling pluripotency, lineage differentiation and fertility. This has prompted many studies to map transcriptional programs during preimplantation development (5–10).

In order to understand regulation of gene expression, precise knowledge of transcriptions start sites (TSSs) is needed. We have used an RNA-seq technology based on the detection of the 5′ ends of transcripts to map active TSSs during the first three days of human preimplantation development (7). *De novo* motif calling using the regions around the detected TSSs led to the identification of multiple significant motifs harboring known transcription factor binding sites (7). Here, we studied a *de novo* motif discovered in these analyses containing a putative binding site for the pleomorphic adenoma gene 1 (PLAG1). *PLAG1* encodes a C2H2 zinc finger transcription factor and an oncogene that was first characterized in pleomorphic adenomas of the salivary gland (11). It belongs to the same protein family as the functionally redundant proto-oncogene PLAG-like 2 (*PLAGL2*) and the imprinted tumor suppressor PLAG-like 1 (*PLAGL1*). Ectopic expression of *PLAG1* and *PLAGL2* resulting from chromosomal translocation events can be found in malignant tumors (12). Associations with cancer have been the focus of most studies on *PLAG* family transcription factors and consequently, less is known about their role in normal physiology. There are no reports on *PLAG* family genes in preimplantation embryo development or pluripotency. By using our human embryo transcriptome data set (13), *Plag1* knockout (KO) mice (14), breeding experiments, single-embryo RNA-seq, and time-lapse analysis of cleavage stage embryo development we show that PLAG1 controls a subset of ZGA genes and is required for normal cleavage stage embryo development.

## RESULTS

### A *De Novo* Motif Containing a PLAG1 Binding Site Is Found in the Promoters of Human ZGA genes

Analysis of the TSSs upregulated during major ZGA in human embryos revealed 13 significant *de novo* DNA motifs, including a 31-bp sequence harboring a known PLAG1 binding site (Figure 1a) (7). This *de novo* PLAG1 motif was found in 74 of the 129 promoters upregulated during ZGA (E = 2.6×10^-646^ by MEME; File S1) and highly similar sites were found in additional 19 promoters (p < 0.0001 by MAST; Figure 1b and File S1). We found that this *de novo* PLAG1 motif was located within a conserved region of an Alu element in close vicinity to an RNA polymerase III promoter A-box. The motif also partially overlapped with the PRD-like transcription factor *de novo* motif that we identified in our previous study as a putative controller of human ZGA (Figures 1c; File S1) (7). Alu elements are primate-specific retrotransposable short interspersed nuclear elements (SINEs) that evolved from a duplication of the 7SL RNA gene (15) and their evolutionary counterparts in rodents are the B1 elements (16). Interestingly, the segment of the Alu element containing the *de novo* PLAG1 motif was well conserved in the rodent B1 element (Figure 1c). Human and mouse PLAG1 are 94% similar by amino acid sequence and their DNA binding domains are identical (Figure S1). The putative PLAG1 binding site within the *de novo* motif in Alu and B1 elements was nearly identical to the PLAG1 consensus site (17) (Figure 1c). Collectively, these data suggest that PLAG1 could be involved in human and mouse ZGA through binding to a conserved motif within Alu and B1 elements. If so, PLAG1 deficiency could lead to disrupted embryonic development and reduced fertility.

**Figure 1.**
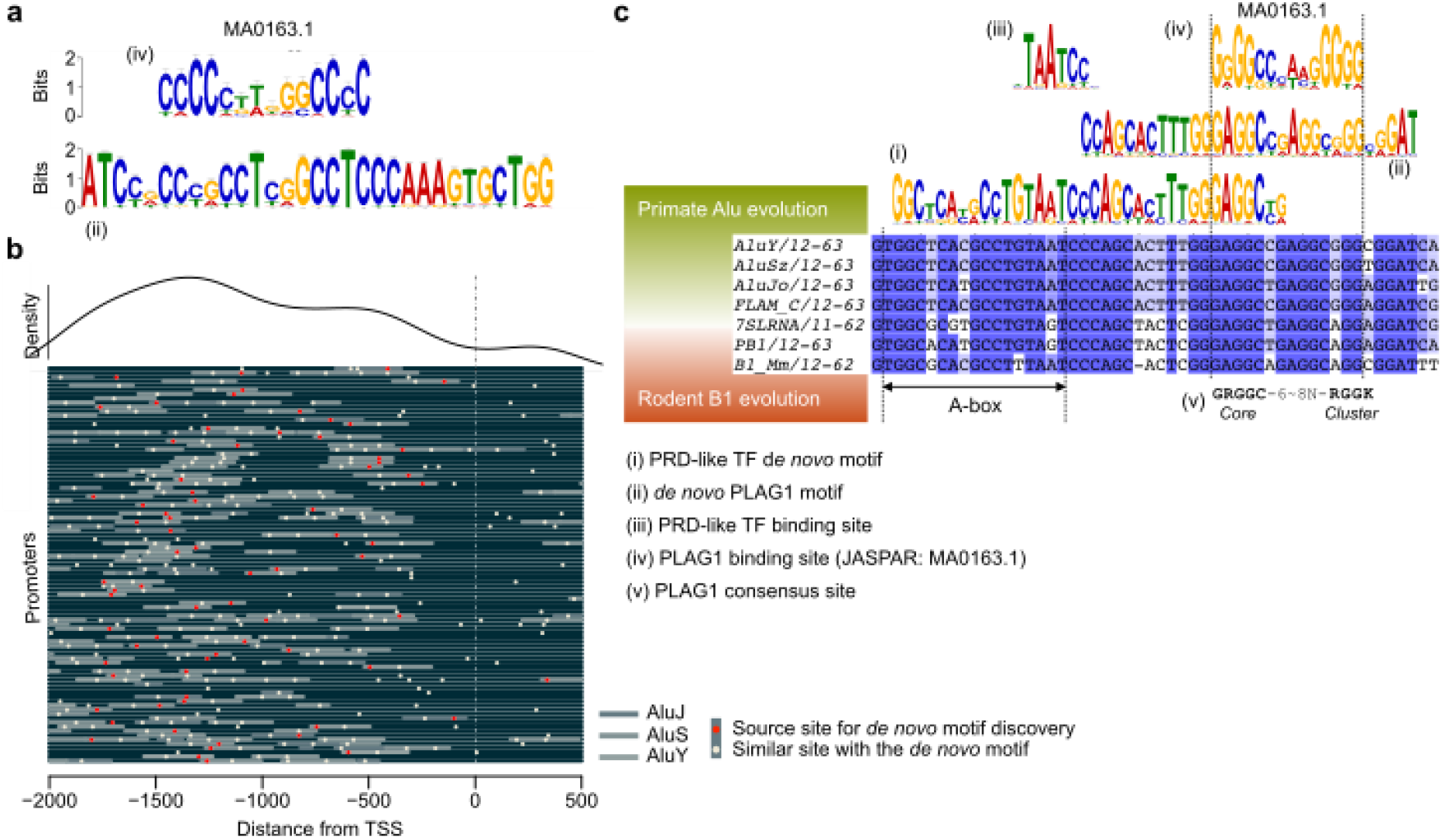
A *De Novo* DNA Motif Containing a PLAG1 Binding Site Is Found in the Promoters of Human ZGA Genes. **(a)** Comparison of (iv) the JASPAR PLAG1 binding site (MA0163.1) and (ii) the *de novo* PLAG1 motif. **(b)** Location of source sites for the *de novo* PLAG1 motifs (red dots) and similar sites (yellow dots) in the promoters of human ZGA genes. Promoters of the 93 genes containing the motifs are stacked and aligned ranging from 2,000 upstream to 500 downstream bases around transcription start site (TSS, dashed line). The genes are listed in File S1. Locations of AluJ, AluS and AluY elements are highlighted in grey. The curve on top of the figure illustrates the density of the *de novo* PLAG1 motif along the promoter. **(c)** Comparisons of (i) the PRD-like transcription factor *de novo* motif, (ii) the *de novo* PLAG1 motif, (iii) the known PDR-like transcription factor binding site, (iv) the known PLAG1 binding site (JASPAR), and (v) the reported consensus PLAG1 binding site (17) with human and mouse short interspersed nuclear elements (SINEs). The consensus motif (v) is described by IUPAC notation in which R is a G or A, and K is G or T. The internal RNA polymerase III promoter, A-box, is indicated. The *de novo* motifs and binding motifs are reverse-complemented. Sequences of AluY, AluSz, AluJo, FLAM_C, 7SLRNA, PB1 and B1_Mm were extracted from the Dfam database of repetitive DNA elements. ZGA, zygotic genome activation; TF, transcription factor.

### *Plag1* Deficiency Affects Reproductive Success But Not Ovarian Or Uterine Function

To study the role of PLAG1 in fertility, *Plag1* KO mice were obtained. The phenotype originally described by Hensen *et al*. was on a Swiss Webster background, while the animals we obtained were backcrossed to the CD-1 strain. Therefore, we started by characterizing the phenotype in our colony. Pups from heterozygous intercrosses (HET×HET) did not significantly deviate from the expected Mendelian distribution (16 litters, 164 pups), although 43% fewer KO pups were born (N = 27) than expected (N = 47) (Figure 2a). Weights recorded at weaning confirmed the earlier reported growth retardation: KO pups were 41% smaller than wildtype (WT) (Figure 2b) (14). Litter frequency over a three-month continuous breeding period did not differ between HET×HET crosses and pairs consisting of a KO female and a HET male (Figure 2c). However, when KO males were crossed with HET females, two of the three pairs did not manage to maintain the approximate one litter-per-month rate (Figure 2c). When we crossed KO mice, litter frequency was significantly reduced compared to HET intercrosses: the three KO×KO breeding pairs produced only two litters in total during the entire three-month test period (Figure 2c).

**Figure 2.**
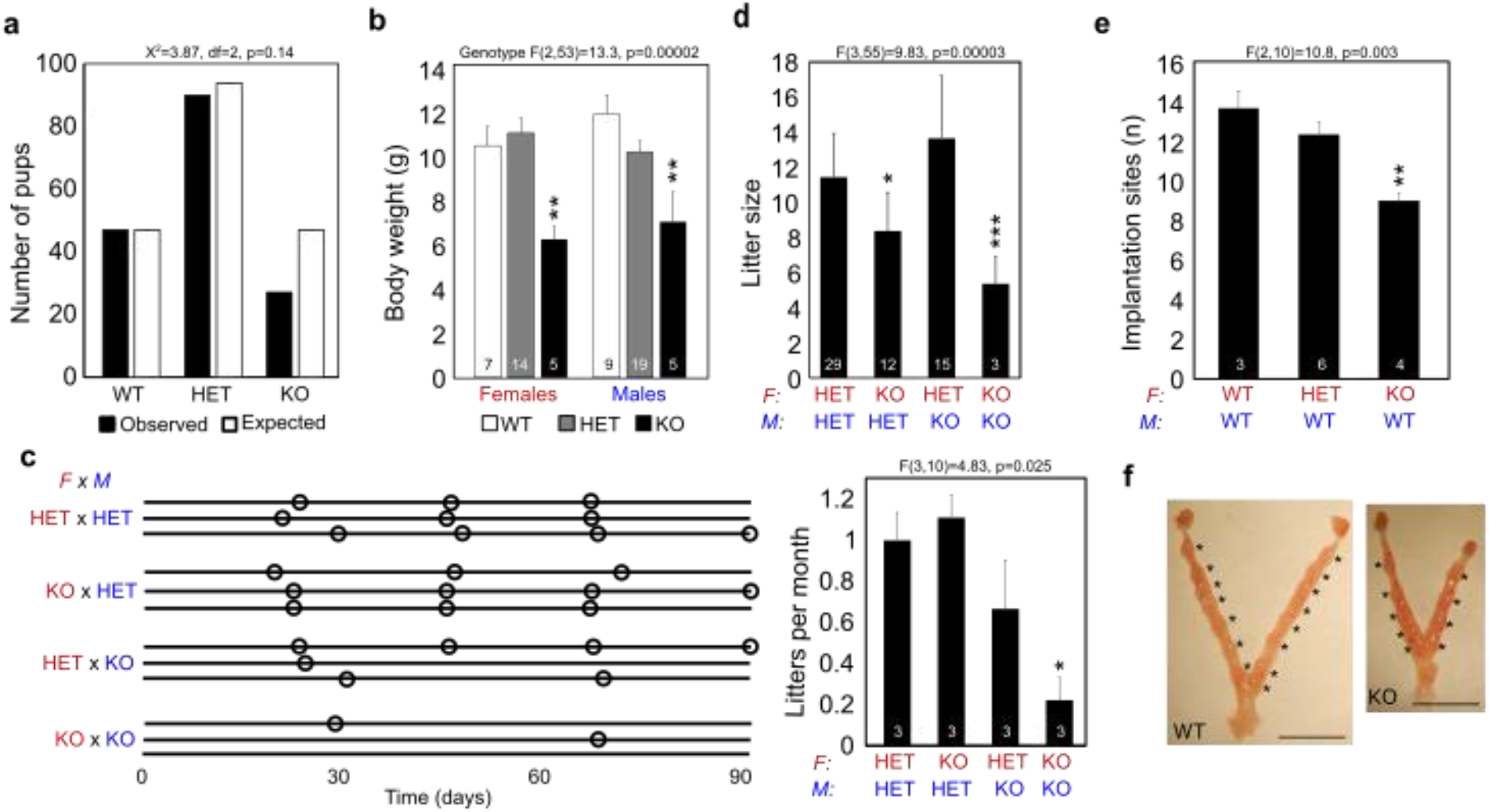
PLAG1 Deficiency Affects Growth and Reproduction in Mice. **(a)** Observed and expected numbers of pup genotypes in litters from *Plag1* heterozygote intercrosses. **(b)** Body weights of female and male pups at weaning (3-weeks-old). **(c)** Frequency of litters from breeding pairs of different genotypes maintained in continuous breeding for 90 days. Every breeding pair is represented with a horizontal line and the birth of a litter is illustrated with a circle. Quantification of the data is shown on the right. **(d)** Mean number of pups in litters from different parent genotypes. **(e)** Mean number of implanted embryos on 7.5–8.5 dpc. **(f)** Representative photos of uteri at dissection. Asterisks indicate implanted embryos. Scale bars are 1 cm. The data in b-e are presented as means + SEM and the number of observations is shown inside the columns. Statistical analysis by χ^2^ test (a) or one-way ANOVA followed by Fisher LSD post-hoc test (b–f). *p < 0.05, **p < 0.01, ***p < 0.001. F, female; HET, heterozygote; KO, knockout; M, male; WT, wildtype.

Compared to HET intercrosses, breeding pairs with a KO female and a HET male produced significantly fewer pups per litter (Figure 2d). With reversed parental genotypes (KO males × HET females), litter size was not affected (Figure 2d). Homozygous KO×KO crosses produced the smallest litters (Figure 2d). The significant reduction in litter size of KO mothers was seen as early as 7.5–8.5 days *post coitum* (dpc), when fewer implantation marks were observed in the uterus compared to HET and WT mothers (Figures 2e, f). As reduced litter size could depend on defects in ovaries or uteri in the *Plag1* KO mothers, we characterized these organs. Oocyte yield in response to gonadotropin-induced superovulation did not differ between WT and KO females (Figure S2a). There were no differences in ovarian weights or numbers of follicles and corpora lutea between the genotypes either (Figures S2b-d). All normal uterine structures were present in both genotypes (Figure S2e). Absolute uterine wet weights were smaller in KO females compared to WT but this difference disappeared when the weights were adjusted for body weights (Figure S2f). To investigate possible differences at the transcriptomic level, RNA-seq was performed on RNA extracted from pieces of uterine horn, endometrial mucosa and myometrium (N = 7). Principal component analysis showed separation of the transcriptomes by tissue type but not by genotype (Figure S2g). Taken together, these data show that *Plag1* KO females have no significant defects in their ovaries and uteri and reproduce with normal frequency, but produce significantly fewer pups per litter as compared to HET and WT females.

### Maternal *Plag1* Knockout Leads to Delayed 2-cell Stage Embryo Development, Disrupted Gene Expression, and Ectopic Expression of paternal *Plag1*

We next studied embryo development. Since *Plag1* KO females had significantly smaller litters compare to HET and WT females regardless of the paternal genotype (Figure 2d, e), we focused on the maternal *Plag1* effect and studied embryos derived from *Plag1* KO females crossed with WT males. These breeding pairs produced HET embryos that lack the maternal *Plag1* allele, and will hereafter be referred to as *matPlag1KO* embryos.

Preimplantation development of 53 WT and 75 *matPlag1KO* embryos was analyzed by time-lapse microscopy from zygote to late morula during three independent imaging sessions, each of which contained both genotypes. Each embryo was followed individually and the time spent in different developmental stages was recorded manually from the composed videos (Videos S1 and S2). We discovered that *matPlag1KO* embryos spent significantly more time in the 2c stage compared to WT embryos (p = 9.99×10^-8^) (Figure 3a). When 50% of WT embryos had already proceeded to the 4c stage, 100% of *matPlag1KO* embryos were still arrested at the 2c stage (Figure 3a, arrow). The time spent in the zygote, 2c and 4c stages was 22.1 ± 0.5 h, 26.1 ± 0.4 h and 14.8 ± 0.3 h (means ± SEM) in the *matPlag1KO* embryos, compared to 21.1 ± 0.3 h, 23.1 ± 0.6 h and 14.3 ± 0.4 h in WT, respectively. On average, *matPlag1KO* embryos spent 3 h longer in 2c stage compared to WT. The survival of the embryos (zygote to morula) did not differ between genotypes: out of the 89 WT zygotes imaged, 53 (60%) developed to a morula, whereas of the 103 *matPlag1KO* zygotes imaged 75 (73%) developed.

**Figure 3.**
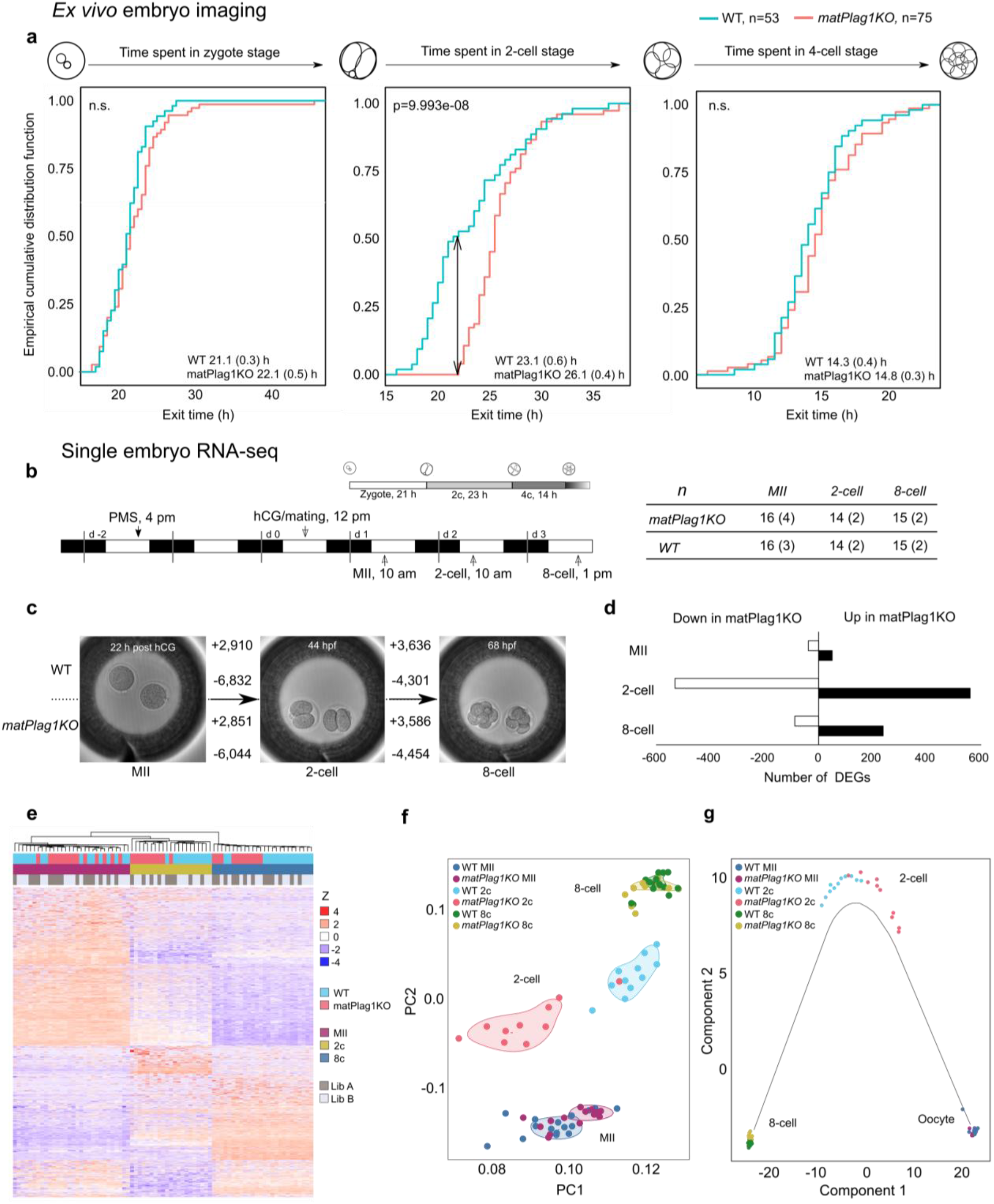
Maternal *Plag1* Knockout Leads to Delayed 2-cell stage Embryo Development and Disrupted Embryonic Gene Expression. **(a)** WT and *matPlag1KO* zygotes were collected for *ex vivo* imaging through time-lapse microscopy and cleavage stage developmental timing was recorded manually for each embryo. Plots show the empirical cumulative distribution function (y-axis, 0–1) of embryos that have exited the corresponding developmental stage at each time point (x-axis, hours). Significance between WT (N = 53) and *matPlag1KO* (N = 75) embryo developmental timing was tested using the Kolmogorov-Smirnov test and p-values are shown in the plots. Average time spent in the stage (SEM) is also shown. **(b)** WT and *matPlag1KO* MII oocytes, 2-cell stage and 8-cell stage embryos were collected for single-embryo RNA-seq. The timing of hormonal treatments, WT embryo development and embryo collection time points are shown together with the numbers of embryos sequenced (numbers of donor females in brackets). **(c)** Number of differentially expressed genes from one developmental stage to another (horizontal arrows) in *matPlag1KO* and WT embryos. **(d)** Number of differentially expressed genes between *matPlag1KO* and WT embryos at the MII oocyte, 2c and 8c stage. **(e-g)** Heatmap (e), principal component analysis (f), and pseudotime (cell trajectory) analysis (g) based on all differentially expressed genes in the libraries. 2c, 2-cell stage; 8c, 8-cell stage; DEG, differentially expressed gene; hCG, human chorionic gonadotropin; KO, knockout; PC, principal component; WT, wildtype.

Next, MII oocytes, 2c embryos and 8c embryos were collected for single-embryo RNA-seq (Figure 3b). In total, we analyzed 45 WT and 45 *matPlag1KO* oocytes and embryos that were collected at three separate time points with both genotypes collected in parallel in every session (Figure 3b). Spike-in RNA was used for data normalization to correct for the large general changes in cellular RNA content during these developmental stages (Figure S3). The majority of differentially expressed genes (DEGs) were downregulated in both genotypes from oocyte to 2c stage when maternal RNA degradation is known to take place (Figure 3c). In that transition, there were 6,832 and 6,044 DEGs downregulated in WT and *matPlag1KO* embryos compared to 2,910 and 2,851 being upregulated, respectively (Figure 3c). More DEGs were upregulated in the 2c–8c transition, when the zygotic genome becomes fully active (3,636 in WT and 3,586 in *matPlag1KO*) (Figure 3c).

The largest difference between WT and *matPlag1KO* embryos was found at the 2c stage (Figure 3d). At this stage, *matPlag1KO* embryos had 530 downregulated and 559 upregulated DEGs compared to WT embryos, suggesting a dysregulation of approximately 11% of all genes regulated during oocyte-to-2c stage transition (Figure 3c,d). This difference was clearly visible in clustering analyses based on all differentially regulated genes in the full dataset. A heatmap clustered the embryos into three primary groups by developmental stage (oocyte, 2c, 8c) and by genotype at the 2c stage (Figure 3e). Principal component analysis likewise separated the embryos primarily by developmental stage and by genotype at the 2c stage (Figure 3f). Finally, cell trajectory (pseudotime) analysis yielded a similar separation at 2c stage, and further suggested that the transcriptional program in the *matPlag1KO* embryos lagged behind the WT at this stage (Figure 3g).

The expression pattern of the DEGs in 2c *matPlag1KO* embryos was studied by plotting their mean expression levels from oocyte to the 8c stage. DEGs upregulated in *matPlag1KO* embryos compared to WT (N = 559) were actually maternally loaded transcripts whose degradation was delayed in the *matPlag1KO* embryos (Figure 4a). Similarly, the DEGs downregulated in *matPlag1KO* embryos at the 2c stage (N = 530) were in fact genes whose upregulation was delayed compared to WT (Figure 4a). We hereafter refer to these two sets of genes as “delayed-degradation” and “delayed-activation” genes. By the 8c stage, the expression levels of the delayed-activation and delayed-degradation genes had reached the WT levels, suggesting that the transcriptional dysregulation caused by the lack of a maternal *Plag1* allele was restored (Figure 4a). To understand the connection of this disrupted gene regulation to *Plag1*, we plotted the expression pattern of *Plag1* by developmental stage in the embryos. *Plag1* transcripts were present in WT oocytes but absent in KO oocytes, as expected (Figure 4b). WT embryos had completely degraded the *Plag1* transcripts by the 2c stage and we did not detect expression in the 8c embryos either, suggesting that *Plag1* is normally not expressed from the zygotic genome (Figure 4b). Interestingly, in clear contrast to WT embryos, *matPlag1KO* embryos expressed *Plag1* in the 2c stage. As these embryos lack maternal *Plag1* allele, the expression must stem from activation of the paternal allele. This ectopic expression of *Plag1* in the 2c stage *matPlag1KO* embryos returned to baseline by 8c stage (Figure 4b), when also the delayed-activation and delayed-degradation genes had caught up with the WT expression levels. This may suggest that the transcriptomic program was rescued by paternal *Plag1* expression.

**Figure 4.**
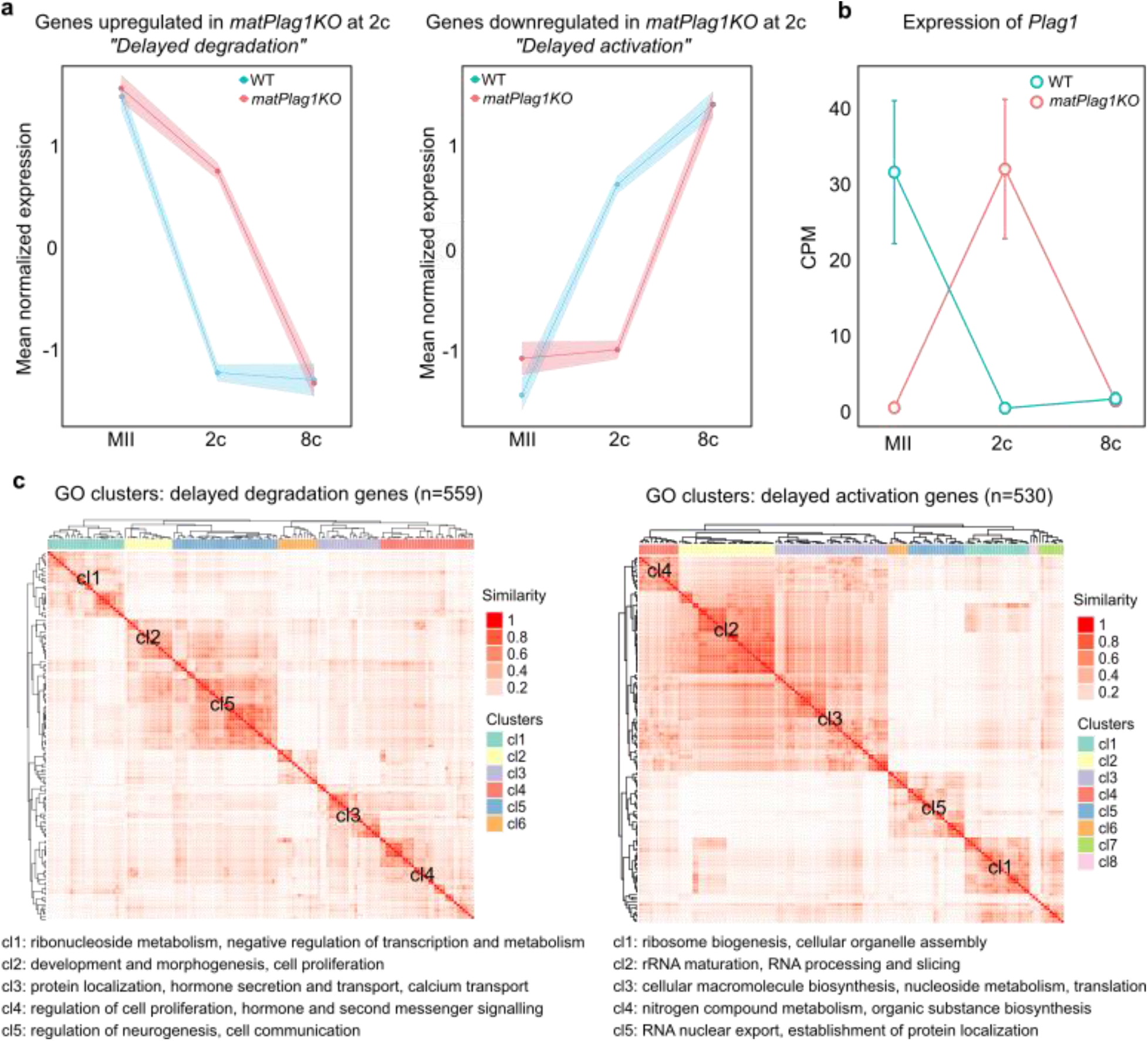
*MatPlaglKO* 2-Cell Embryos Have Delayed Regulation of Two Functionally Differing Sets of Genes and They Ectopically Express *Plag1* from the Paternal Allele. **(a)** Mean normalized expression (Z) pattern of the 559 genes that are significantly upregulated (“delayed-degradation”) and the 503 genes that are significantly downregulated (“delayed-activation”) in *matPlag1KO* embryos compared to WT embryos at the 2c stage. **(b)** Normalized expression of *Plag1* in the MII oocyte, 2c and 8c stages. **(c)** Heatmaps displaying semantic similarity among the top-150 significantly enriched GO terms associated to the delayed-degradation (left) and delayed-activation (right) genes. Five largest clusters are depicted (cl1–cl5), and common denominators among GO terms belonging to these clusters are shown. The full GO lists are provided in File S2. 2c, 2-cell stage; 8c, 8-cell stage; CPM, counts per million; GO, gene ontology; KO knockout; WT, wildtype.

ZGA takes place with the help of maternally provided factors deposited to the oocytes during folliculogenesis. Our data show that *Plag1* is a maternally provided transcription factor in the mouse (Figure 4b), and analysis of two independent human datasets showed that *PLAG1* is also maternally provided to human oocytes (Figure S4). In addition, we stained mouse ovary sections for X-gal, the marker of *Plag1* expression in the KO mice, and found positive staining in the nuclei of oocytes within growing follicles, further demonstrating that *Plag1* is a maternally provided factor (Figure S5).

The functions of the delayed-activation and delayed-degradation genes were studied via gene ontology (GO) annotations. The delayed-activation genes clustered into categories relevant to ribosome biogenesis, RNA processing, and translation (Figure 4c; File S2), whereas the delayed-degradation genes showed less clear clustering representing diverse GO categories ranging from neurogenesis to negative regulation of transcription (Figure 4c, File S2). These GO categories were compared to those associated with normal mouse embryo development (File S3). The delayed-activation GOs showed higher similarity to genes typically upregulated during 2c–8c transition in the WT [best-match average (BMA) 0.792] than to those downregulated (BMA 0.531), whereas the delayed-degradation GOs showed higher similarity to genes normally downregulated between 2c–8c stages (BMA 0.714) than to those upregulated (BMA 0.445). These data confirmed that embryos that lacked the maternal *Plag1* allele were characterized by delayed transition through the 2c stage due to specific sets of genes being dysregulated compared to WT embryos.

### The Delayed-Activation Genes Overlap with Human ZGA Genes and Have Enriched PLAG1 *De Novo* Motif in Their Promoters

We next wanted to compare our current mouse embryo data with our earlier human embryo transcriptome data (13). To overcome species differences, we converted the genes to orthologues and compared both gene and protein families. The gene expression changes during major ZGA in humans (4c–8c transition) and WT mice (2c–8c transition) were highly similar in general between our datasets (Figure S6). A gene set analysis showed that the TSSs upregulated during human major ZGA were significantly enriched among the mouse delayed-activation genes (Figure 5a). The same result was obtained when using the more conservative χ^2^ test that is based on significant DEGs only; a significant overlap was found between the human major ZGA genes with the *matPlag1KO* delayed-activation genes but not with the delayed-degradation genes (Figure 5b). Comparison of protein families yielded similar results (Figure 5c).

**Figure 5.**
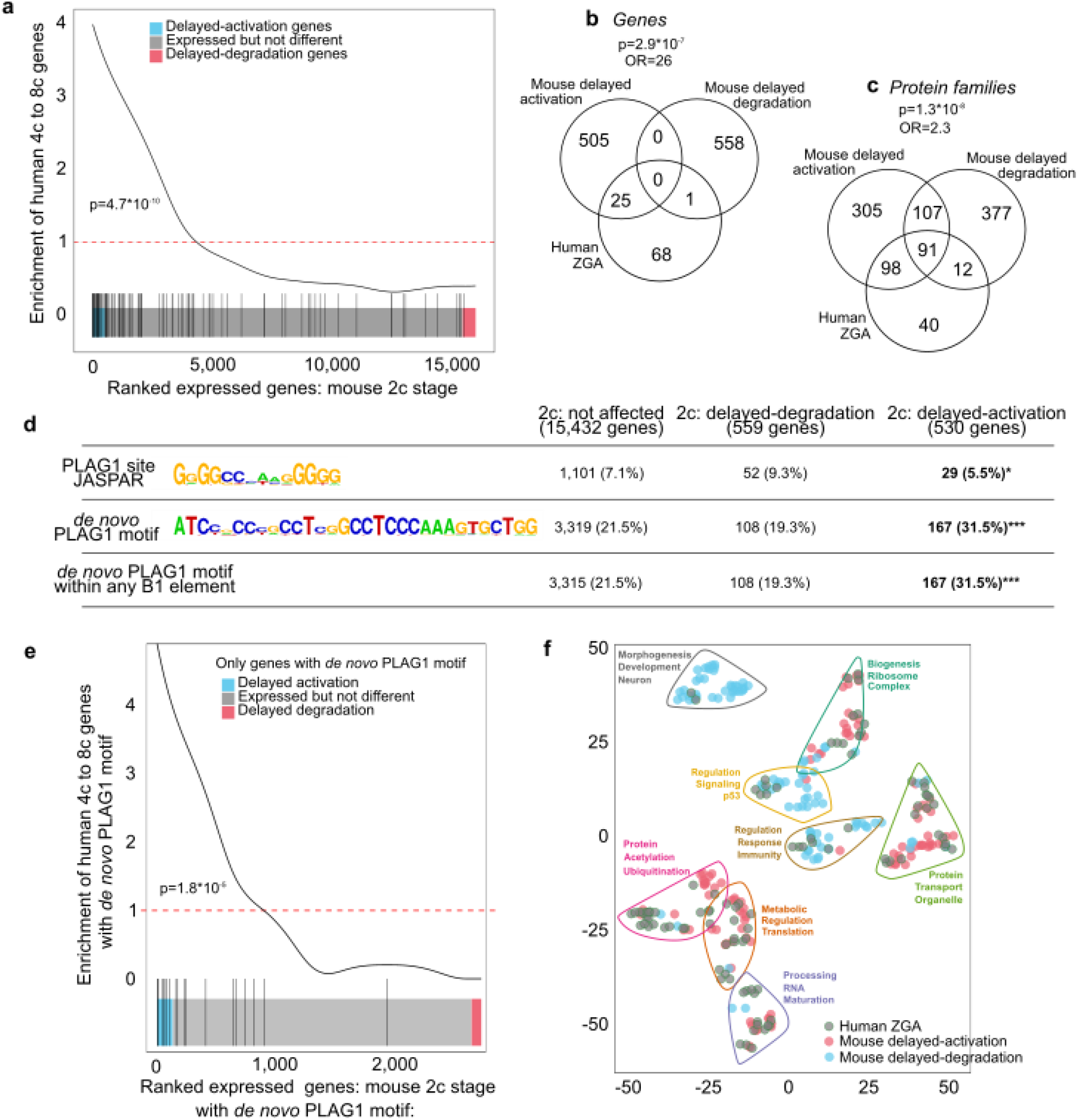
*MatPlag1KO* Delayed-Activation Genes Overlap with Human ZGA Genes, Have Enriched PLAG1 *De Novo* Motif in Their Promoters, and Associate with Protein Synthesis. **(a)** Gene set enrichment analysis comparing genes that are expressed in mouse embryos (WT and *matPlag1KO*) at the 2c stage (x-axis) with the genes upregulated during human ZGA (4c–8c transition) (y-axis). The black vertical lines shows the location of orthologous human genes among the ranked mouse genes, and the curve depicts the enrichment. Red dotted line indicates “no enrichment” level. Significance was tested with the GeneSet test function. **(b-c)** Venn diagram showing the overlap between delayed-activation and delayed-degradation genes (b) and proteins (c) with the human ZGA genes. **(d)** Presence of PLAG1 binding sites (JASPAR), *de novo* PLAG1 motifs, and *de novo* PLAG1 motifs within B1 elements in the promoters of genes expressed in mouse 2c embryos. Total number of genes in different categories as well as number of unique genes with the motif within -2,000 to +500 of their TSS are shown. Enrichment over not affected genes was analyzed with Fisher’s exact test. ***p < 0.001. **e)** Gene set enrichment analysis comparing human and mouse genes that contain at least one *de novo* PLAG1 motif in their promoters. Significance was tested with the GeneSet test function. **(f)** t-SNE plot demonstrating similarity among the top-100 GO categories associated to human ZGA, mouse delayed-activation and mouse delayed-degradation genes that contain at least one *de novo* PLAG1 motif in their promoters (within -2,000 to +500 bp of TSS). The eight largest clusters with the most common words within the clusters are shown. 2c, 2-cell stage; 4c, 4-cell stage; 8c, 8-cell stage; GO, gene ontology; KO, knockout; OR, odds ratio; TSS, transcription start site; WT, wildtype.

We then studied the occurrence of PLAG1 binding sites (MA0163.1 in JASPAR) and *de novo* PLAG1 motifs in the affected mouse gene promoters within -2,000 bp and +500 bp of their TSS. One or more JASPAR PLAG1 binding sites were found in less than 10% of the promoters (Figure 5d). Surprisingly, even fewer (5.5%) delayed-activation genes contained the site than the genes that were not affected by *Plag1* KO (7.1%) (Figure 5d). However, the results were the opposite when we used our *de novo* PLAG1 motif instead of the JASPAR PLAG1 site. A significantly higher proportion of the delayed-activation promoters contained at least one *de novo* PLAG1 motif (31.5%) compared to “expressed but not affected” (21.5%) and delayed-degradation (19.3%) genes (Figure 5d; File S4). We also studied the occurrence of the *de novo* PLAG1 motifs located within B1 elements and obtained virtually identical results, showing that the vast majority of the *de novo* PLAG1 motifs reside within B1 elements (Figure 5d). In addition to occurrence (present or not), we also considered the frequency (how many) of the *de novo* PLAG1 motifs in the mouse promoters. A significantly higher frequency was found in the promoters of delayed-activation genes compared to delayed-degradation genes (Figure S7a,b).

Finally, we compared the homologous mouse delayed-activation and human ZGA genes that contained at least one *de novo* PLAG1 motif and found a significant overlap (Figure 5e). We GO-annotated the three gene groups, *i.e*. human ZGA, mouse delayed-activation and mouse-delayed degradation genes. Hierarchical clustering revealed that human ZGA and mouse delayed-activation genes clustered together to categories representing ribosomes, protein transport, translation, RNA processing, and protein metabolism, whereas mouse delayed-degradation genes represented neurogenesis, p53 signaling and immune responses showing little overlap with the human ZGA clusters (Figure 5f). These data suggest that mouse and human embryos have a functionally conserved set of genes that are activated during ZGA and contain enriched PLAG1 binding sites in their promoters (File S4).

## DISCUSSION

In the present study, we have discovered a new role for the oncogene *PLAG1* in the regulation of ZGA. We identified a *de novo* assembled motif containing a PLAG1 binding site among the promoters upregulated during ZGA in human embryos, and showed that the lack of maternally loaded *Plag1* in mouse oocytes lead to a significant delay in ZGA on a transcriptional level with consequences for the timing of cleavage-stage development. Restoration of gene expression levels and embryo developmental speed coincided with ectopic expression of *Plag1* from the paternal allele. These data imply an essential functional role for PLAG1 in the regulation of ZGA. Our data further propose that PLAG1 target genes have roles in central cellular processes that relate to ribosomes, RNA and protein metabolism, which undoubtedly have an essential function during early embryo growth.

Our studies on the *Plag1* KO mice, backcrossed to the CD-1 strain, confirmed their growth restriction and subfertility originally observed in the Swiss strain (14). We further expanded the observations by Hensen et al. (2004) by showing that KO females produced small litters regardless of the paternal genotype, which implies that this phenotype was caused by the lack of a *Plag1* allele in the mother. We did not find any significant differences in histology or function of reproductive organs between KO and WT females, except for smaller uterine wet weights in the KO females. However, the body weight adjusted uterine weights did not differ between the genotypes. It could be still argued that smaller uterine size imposes spatial constraints on the number of embryos that the uterus can accommodate. However, this does not seem to be the case. Various mutant mouse strains show growth restriction or dwarfism, but still produce normal litter sizes compared to WT controls. Examples include *Ccnd1* KO (18) and paternal *Mest* KO (19). Furthermore, “uterine crowding” has not been found to affect fetal viability in rabbits (20) or mice (21). Lastly, litter size was even more affected when KO females were crossed to KO males than when they were crossed to HET males, despite the fact that the uteri of the KO females were of similar size in both crosses. We therefore conclude that the reduced litter size observed in *Plag1* KO females is unrelated to their smaller uteri.

Our data suggest that HET embryos derived from KO females by crossing them with WT, i.e. *matPlag1KO* embryos, are not only different from WT but also from HET embryos produced by reversing the parental genotypes. We show that *matPlag1KO* embryos that lack the *Plag1* transcript only during the first hours following fertilization, spent significantly longer time in the 2c stage compared to WT embryos. The 2c stage is the developmental stage when the major ZGA takes place in the mouse, and failure to activate transcription leads to developmental arrest (3). Embryo development can be affected by single genes. For example, knockdown of the pluripotency factor *Lin28* in mouse embryos leads to arrest at the 2c–4c stage (22), and KO of the maternal-effect gene *Mater* to arrested embryonic development at the 2c stage (23). ZGA is dependent on maternally provided factors, *i.e.*, transcripts and proteins that are deposited into the oocyte during folliculogenesis. We showed that *Plag1* is a maternally provided factor both in mice and humans. The transcript could be detected in mature oocytes by RNA-seq in both species. Although there is a lack of specific antibodies against mouse PLAG1, we were able to show PLAG1 protein expression in growing mouse oocytes indirectly via X-gal staining since the *Plag1* coding sequence is replaced by that of *lacZ* in the mutant mice (14). The mutant locus is flanked by *Plag1* untranslated regions, giving the *lacZ* transcript the same translational control as the *Plag1* transcript would have. Taken together, we show that *Plag1* is a maternally provided factor and its absence in oocytes leads to delayed 2c stage development, suggesting that *Plag1* is required for normal cleavage-stage embryo development.

Interestingly, at the 2c stage, when the development of every studied *matPlag1KO* embryo was delayed compared to WT, these embryos started expressing *Plag1* from the paternal allele. This upregulation was followed by regaining the normal developmental pace as well as normal transcriptomic program by the 8c stage, suggesting that the ectopic paternal expression of *Plag1* rescued the embryo phenotype. Although these data argue that *Plag1* is essential for embryo development at the 2c stage, we noted that even homozygous *Plag1* KO intercrosses occasionally produced litters. We hypothesize that *Plagl2*, the *Plag1*-family member with redundant functions to *Plag1* that is also maternally provided, might rescue embryo development in some cases. Testing this hypothesis would require generation of *Plag1/Plagl2* double KO mice, which is impossible due to the severe phenotype of *Plagl2* KO mice; KO pups die shortly after birth to starvation due to their inability to absorb chylomicrons (24).

Although *matPlag1KO* embryos regained a normal developmental pace after exiting the 2c stage, their developmental success was not the same as that of WT embryos, as evidenced by the reduced number of implantation sites and smaller litter size. Our embryo time-lapse data do show that *matPlag1KO* embryos developed to the blastocyst stage with equal efficiency as WT embryos. Since KO uteri already showed less implantation marks at 7.5–8.5 dpc, the embryo loss must occur sometime between 4.5 and 7.5 dpc, *i.e.*, when the blastocyst normally implants or shortly thereafter. Interestingly, human embryos can be scored after *in vitro* fertilization through morphokinetic measurements, and the time an embryo spends in the 2c and 4c stages is a significant determinant of developmental potential (25). Following this, we hypothesize that the delayed 2c stage development and associated dysregulation of over 1,000 genes in *matPlag1KO* embryos could have adverse effects on the overall developmental potential of the blastocysts. In addition, synchronous preparation of both the embryo and endometrium for implantation is a prerequisite for successful implantation during the window of receptivity. Therefore, a simple delay in preimplantation development could contribute to some blastocysts missing this critical window (26).

The *de novo* PLAG1 motif in humans frequently localized within Alu elements in the promoters of the ZGA genes. Alu elements are transposable elements ubiquitously present in primate genomes with involvement in gene regulation through various mechanisms (27). The counterparts of Alu elements in the mouse genome are B1 elements (16). Mouse B1 and primate Alu elements have split from a common ancestor, the 7SL RNA gene, over 80 million years ago and have evolved and retrotransposed independently since (15). Despite independent evolution, the densities of these elements in promoters of orthologous genes in humans and mice are surprisingly correlated (28, 29). It is interesting that the component of the Alu element containing the PLAG1 binding site was well-conserved in the mouse B1 element, suggesting positive evolutionary selection for this sequence. It has also been shown that genes containing Alu and B1 elements are highly enriched in promoters that are activated during ZGA (30). We confirm this in our datasets, showing that both human and mouse ZGA gene promoters are enriched for Alu/B1 elements (containing the *de novo* PLAG1 motif) and the associated genes relate to ribosomes in both species. Collectively, these findings may suggest that Alu and B1 play a role in ZGA in humans and mice by attracting transcription factors such as PLAG1 to gene promoters. Our data further showed that the PLAG1 binding site as reported in the JASPAR database alone did not show enrichment in the promoters of delayed-activation genes and that the site in general was more rare than the *de novo* PLAG1 motif. This suggested that PLAG1 works in concert with other transcription factors and co-regulators and therefore the short binding site does not accurately reflect the DNA sequence required for PLAG1-stimulated gene expression. Co-factors and binding motif preferences of different PLAG1 protein complexes is a largely unexplored area that should be a focus of follow-up studies.

Functional annotation categories associated with delayed-activation genes in the *matPlag1KO* embryos were mainly related to ribosome biogenesis, maturation and function. Even when we restricted the analysis to those genes that contained the *de novo* PLAG1 motif, GO analyses suggested roles in ribosome biogenesis and RNA and protein metabolism, both in mice and humans. It has been shown that Alu elements are enriched in promoters of genes involved in ribosome biogenesis, protein biosynthesis and RNA metabolism (29). It is a plausible hypothesis that Alu elements in the promoters of ribosomal-function genes attract PLAG1 that then regulates the associated gene. Many oncogenes have effects on ribosome biogenesis, which enables cancerous cells to increase protein synthesis and grow rapidly (31). Although *PLAG1* is an oncogene, its potential role in ribosome biogenesis and protein synthesis has not been investigated. One of the most striking phenotypes of the *Plag1* KO mice is their small size (14). In addition, *PLAG1* polymorphisms associate with body size and growth in farm animals and humans (12, 32–35). Based on our data, we present a hypothesis that PLAG1-associated growth phenotypes, such as growth retardation in the KO mice, results from modulation of protein synthesis that affects cell size and division rate.

We conclude that the lack of maternally provided *Plag1* leads to a delay in ZGA manifested as prolonged 2c stage development, which is rescued by ectopic paternal *Plag1* expression. This delay and associated dysregulation of genes needed for ribosome biogenesis, RNA and protein metabolism could lead to reduced embryo competence for implantation, explaining the reduced litter size in KO mothers. We further propose that the effect of PLAG1 on ZGA genes arose through retrotransposition of Alu and B1 elements, where the PLAG1 binding sites have been under positive selection. Follow-up studies should focus on a deeper analysis of PLAG1 involvement in protein synthesis, as this is a mechanism that would explain many of the reported biological activities of PLAG1, including tumorigenicity, cell proliferation, and growth.

## Supporting information

## ACKNOWLEDGMENTS

*Plag1* KO mice were generated in the Laboratory for Molecular Oncology of Prof. Wim Van de Ven, Center for Human Genetics, Catholic University of Leuven, Belgium. The authors thank Prof. Wim Van de Ven for access to the *Plag1KO* mouse line and Dr Carol Schuurmans (University of Calgary, Canada) for providing the mice. Stanisław Wawrzyczek is thanked for assistance with X-gal staining and Ingegerd Fransson for RNA-seq library preparation. The morphological phenotype analysis (FENO) core facility and Tarja Schröder are thanked for the preparation of the histological specimens. This study was performed in part at the Preclinical Laboratory (PKL) animal facility and the Live Cell Imaging unit, Department of Biosciences and Nutrition, Karolinska Institutet, Sweden, that are supported by grants from the Knut and Alice Wallenberg Foundation, the Swedish Research Council, the Centre for Innovative Medicine and the Jonasson donation to the School of Technology and Health, Royal Institute of Technology, Sweden. The computations were performed on resources provided by SNIC through Uppsala Multidisciplinary Center for Advanced Computational Science (UPPMAX) under projects b2010037 and snic2017-7-317. The study was supported by the Knut and Alice Wallenberg Foundation (KAW2015.0096) and by a Distinguished Professor Award from Karolinska Institutet to JK. Essential funding was also provided by the Jane & Aatos Erkko Foundation to PD.

## AUTHOR CONTRIBUTIONS

P.D., E.M. and J.K. planned and designed the study. P.D. and E.M. carried out experiments. P.D., A.D. and E.M. analyzed data; A.D. and S.K. designed and carried out bioinformatic analyses; K.K. and E.E. performed RNA-seq analyses; K.M. conducted histomorphometric measurements and embryo morphokinetic analyses; and B.D.G. did X-gal stainings. J.K. and O.H. provided essential resources. P.D., E.M., A.D., B.D.G. and J.K. wrote the paper. All authors commented on the manuscript and have approved the final version.

## DECLARATION OF INTERESTS

The authors declare no competing interests.

## MATERIALS AND METHODS

### Mouse Strain and Husbandry

All experiments were approved by the Swedish Board of Agriculture (#S5-14) and performed in accordance with the ethical licence. *Plag1KO* mice (14), backcrossed to the CD-1 strain, were a kind gift from Prof. Wim Van de Ven (University of Leuven, Belgium) and Dr. Carol Schuurmans (University of Calgary, Canada). The local colony at Karolinska Institutet was established through embryo transfers. HET animals were kept in continuous breeding, and the litters were earmarked, weaned and weighed at the age of 3–4 weeks. Ear punches were used for genotyping with primers Plag1MT_F (5′-CAGTTCCCAGGTGTCCAACAAG-3′), Plag1MT_R (5′-AATGTGAGCGAGTAACAACCCG-3′), Plag1WT_F (5′-CGGAAAGACCATCTGAAGAATCAC-3′), and Plag1WT_R (5′-CGTTCGCAGTGCTCACATTG-3′). The animals were housed in standard conditions (19–21°C, 55% humidity, lights 6:00 am–6:00 pm) with free access to feed (irradiated Global 18% diet 2918; Envigo, Huntingdon, UK) and tap water. In all experiments, animals were sacrificed by cervical dislocation.

### Superovulation and Embryo Imaging

Sexually mature 1–3-months-old *Plag1* KO and WT females were superovulated by i.p. injection of 5 IU pregnant mare serum (Folligon; Intervet, Dublin, Ireland), followed two days later by 5 IU human chorionic gonadotropin (hCG) (Chorulon; Intervet, Dublin, Ireland), and mated with trained WT studs. MII oocytes (no mating) and zygotes were collected from oviducts the following morning. Cumulus cells were removed with hyaluronidase (0.3 mg/ml, Sigma-Aldrich, St.Louis, MO, USA). For imaging, zygotes were placed 2–4 per well into Primo Vision embryo culture dishes (Vitrolife, Göteborg, Sweden) under a Nikon Ti-E spinning disk wide-field microscope with a live-cell imaging incubator. The microscope was programmed to take bright-field images every 30 min with an Andor EM-CCD camera using the 20× objective for a total of 90 h 30 min. The imaging was repeated three times with embryos from 5 KO females (providing 103 embryos) and 6 WT females (providing 89 embryos) with both genotypes present at every session. Cleavage events until the 8c stage were scored manually from the images by a researcher blinded to the genotypes.

### Histological Assessment of Ovaries and Uteri

Ovaries and uteri were collected during superovulation experiments, their weights recorded, and tissues stored in 4% (w/v) paraformaldehyde. Paraffin-embedded tissue was processed to 4-μm hematoxylin-eosin-stained sections and slides digitalized with a Mirax Slide Scanner (Zeiss, Göttingen, Germany). Transverse sections from the middle of the uterine horn were used for histological examination. One section from the middle of the ovary was used for assessment of follicles in different developmental categories (preantral, antral, atretic and corpora lutea) and their number adjusted for the ovary surface area using Pannoramic Viewer software (3DHistech, Budapest, Hungary). Altogether, 22 KO and 13 WT animals were used for assessment of the ovary, and 8 KO and 8 WT for uterus histology.

### X-gal Staining

Ovaries were collected from 3 KO and 3 WT females, aged between 4 and 5 months, fixed in 4% paraformaldehyde, cryosectioned at 12 μm and subjected to X-gal staining as described before (36), with the following modifications: sections were placed in X-gal solution at room temperature immediately without thawing; after overnight incubation in X-gal solution, the tissues were not postfixed; and eosin was used as a counterstain.

### Implantation Study

KO and WT females, aged 2-4 months, were mated with trained WT studs. The uteri were collected 7.5–8.5 dpc and the number of implantation sites counted. The experiment was carried out twice, with a total of 8 KO and 8 WT females.

### Single-Embryo RNA-seq

Zygotes collected from superovulated females were treated with hyaluronidase and placed in KSOM medium (EMD Millipore, Nottingham, UK) under ovoil-100 (Vitrolife) in IVF 4-well plates (Sigma-Aldrich) for culture at 37°C under 5% CO2/95% air. Single embryos or uncultured MII oocytes were picked into 4 μl library lysis buffer containing 5 mM Tris-HCl pH 8.0 (Sigma-Aldrich), 2 mM dNTP mixture (ThermoFisher Scientific, Waltham, MA, USA), 10 mM DTT (Sigma-Aldrich), 0.05% Triton X-100 (Sigma-Aldrich), 400 nM anchored oligo(dT) primer biotin-TTAAGCAGTGGTATCAACGCAGAGTCGAC(T)_29_V where V is a LNA nucleotide (Exiqon, Vedbaek, Denmark), and 4 U RiboLock RNase inhibitor (ThermoFisher). Embryos were picked as shown in Figure 3b on three different occasions, yielding a total of 16 zygotes, 14 2c-stage embryos, and 15 8c-stage embryos for both *matPlag1KO* and WT (Figure 3b). Two separate 46-plex libraries (libA and libB) were made as described previously (37) with the following modifications: barcoded 10 μM template-switching oligonucleotides were added prior to reverse transcriptase, ERCC spike-in Mix A was diluted 15,300-fold with clean water, and 1 μl was taken per library reverse transcriptase master mix. Twenty cycles of PCR were used for the first round of amplification and ten additional cycles for the second round to introduce Illumina-compatible universal sequences. Both libraries contained all developmental stages and genotypes.

### Uterus RNA-seq

Uterine horns from 8 *Plag1KO* and 8 WT females used in the superovulation experiments were collected for RNA-seq. At dissection, uterine horns were cut in half longitudinally, and the endometrial side was gently scraped with a scalpel to separate the mucosa from myometrium, and the samples were stored in RNAlater (Ambion, Foster City, CA, USA). RNA was extracted with the RNeasy Mini kit (Qiagen, Hilden, Germany) and quality measured with the Agilent 2100 BioAnalyzer (Agilent Technologies, Santa Clara, CA, USA). Ten nanograms of high-quality RNA (RIN>8) was used for RNA-seq. RNA-seq was performed according to the modified STRT protocol (37).

### RNA-seq Data Analysis

The analysis was performed as described previously (37). Briefly, the reads were filtered, samples de-multiplexed, UMIs joined, reads trimmed and mapped to the reference mouse genome mm9 by TopHat (38). The resulting bam files were converted to tag directories employing Homer (39) and were subsequently used to estimate the reads in all annotated genes. Annotations were in GTF file format retrieved from UCSC and were concatenated to a GTF file with the ERCC annotations. Gene counts were then imported to R (40) and libraries with a median gene expression of log2 counts per million (cpm) under 0 were excluded from further analysis. Cell libraries, excluding the ERCC spike-in counts, were normalized with EdgeR (41) using the TMM normalization method. The ERCC counts were used for normalization between the various embryonic cell stages by scaling the library sizes. EdgeR was also employed for the subsequent differential gene expression analysis, which was performed on genes that had 1 cpm in at least five or more samples and the rest of the genes were filtered out. After removal of low-expressed and unexpressed genes, the gene counts were renormalized. Principal component analysis was performed in R by using the genes that were significant in any of the comparisons between the genotypes. Heatmaps were plotted on TMM-normalized counts exported from EdgeR and gene expression was standardized across all samples (mean = 0 and SD = 1). Samples and genes were clustered using hierarchical clustering in R and plotted employing the ComplexHeatmap library (42). The same gene set was used for the cell trajectory (pseudotime) analysis by the monocle package (43). Gene ontology analysis was carried out in R with the topGO library. To identify enriched GO terms, the classic algorithm and Fisher statistic were used and analysis was carried out on up- and downregulated genes separately. Semantic similarity between the GO terms was calculated using the Wang algorithm in the GOSemSim bioconductor package (44). The result is given as the best-match average (BMA) score that ranges from 0 to 1. Gene set enrichment analysis was conducted to test whether the mouse genes homologous to the human genes regulated between the 4c and 8c stage were also regulated in mouse development. To calculate the p values, the geneSetTest function from the *limma* package (45) was used. The moving average of the enrichment was calculated with the tricubeMovingAverage function and plotted with ggplot2. Homologene from NCBI was used to convert the human genes to the homologous mouse genes. The significance in overlap of the human genes with the mouse genes regulated in the KO at the 2c stage was calculated with the Fisher test in R. Genes were also converted to protein families using the bioconductor libraries for genome-wide annotation for human and mouse (org.Hs.eg.db/org.Mm.eg.db). For genes that had more than one protein family annotated to them, only one of the protein families was used in order not to inflate the number of overlapping or non-overlapping families between the different gene groups.

### Promoter Analyses

Human embryo promoter analysis was performed as previously described (13). Briefly, the *de novo* motif was compared with known motifs by TomTom (46). We applied MEME (47) for motif analysis within the upregulated promoters, and identified sequences similar to the PLAG1 motif (MA0163.1 in JASPAR) (48, 49) using MAST (50). The location of Alu elements within the promoters was based on the RepeatMasker track in the UCSC Genome Browser. Human and mouse SINE repetitive elements DF0000002 (AluY), DF0000051 (AluSz), DF0000034 (AluJo), DF0000144 (FLAM_C), DF0000016 (7SLRNA), DF0003101 (PB1) and DF0001733 (B1_Mm) were retrieved from the Dfam database (51) and aligned and highlighted according to the percent identity by JalView2. Mouse embryo promoter analysis was carried out with Homer (39). The hits of the repetitive elements and the motifs were imported into R. Repetitive elements that fell within the promoter regions (TSS -2,000 bp to +500bp) were kept and the distance to the nearest TSS calculated for the repeats and the *de novo* PLAG1 motifs and plotted using ggplot2. Enrichment was analyzed using Fisher’s exact test.

### Statistical Analysis

Continuous data were analyzed with Student’s *t*-test, one-way ANOVA (one categorical predictor) or two-way ANOVA (two categorical predictors), followed by Fisher LSD post-hoc test when necessary. Normality was tested with the Shaphiro-Wilks test and homoscedasticity with Levene’s test. Categorical data were analyzed with the χ^2^ test. The exit time of embryos from each developmental stage was plotted as an empirical distribution function (ecdf) using ggplot2. To test the significance between the exit times of the WT and *matPlag1KO* embryos, the Kolmogorov-Smirnov test was used. All analyses were carried out using R. All p values were two-tailed and were considered significant if p < 0.05.

## DATA AVAILABILITY

RNA-seq data have been deposited to Gene Expression Omnibus data repository as a SuperSeries record under the reference GSE111040.

## Notes

#### Summary of Updates

Figures have been rearranged for clarity, and new data comparing mouse and human ZGA genes containing de novo PLAG1 motifs has been added.

## REFERENCES

1. Jukam D, Shariati SAM, & Skotheim JM (2017) Zygotic Genome Activation in Vertebrates. Dev Cell 42(4):316–332.

2. Niakan KK, Han J, Pedersen RA, Simon C, & Pera RA (2012) Human pre-implantation embryo development. Development 139(5):829–841.

3. Warner CM & Versteegh LR (1974) In vivo and in vitro effect of alpha-amanitin on preimplantation mouse embryo RNA polymerase. Nature 248(5450):678–680.

4. Abe KI, et al. (2018) Minor zygotic gene activation is essential for mouse preimplantation development. Proc Natl Acad Sci U S A 115(29):E6780–E6788.

5. De Iaco A, et al. (2017) DUX-family transcription factors regulate zygotic genome activation in placental mammals. Nat Genet 49(6):941–945.

6. Petropoulos S, et al. (2016) Single-Cell RNA-Seq Reveals Lineage and X Chromosome Dynamics in Human Preimplantation Embryos. Cell 165(4):1012–1026.

7. Töhönen V KS, Vesterlund L, Sheikhi M, Antonsson L, Filippini-Cattaneo G, Jaconi M, Johnsson A, Linnarsson S, Hovatta O, Kere J (2017) Transcription Activation Of Early Human Development Suggests DUX4 As An Embryonic Regulator. bioRxiv.

8. Vassena R, et al. (2011) Waves of early transcriptional activation and pluripotency program initiation during human preimplantation development. Development 138(17):3699–3709.

9. Xue Z, et al. (2013) Genetic programs in human and mouse early embryos revealed by single-cell RNA sequencing. Nature 500(7464):593–597.

10. Yan L, et al. (2013) Single-cell RNA-Seq profiling of human preimplantation embryos and embryonic stem cells. Nat Struct Mol Biol 20(9):1131–1139.

11. Kas K, et al. (1997) Promoter swapping between the genes for a novel zinc finger protein and beta-catenin in pleiomorphic adenomas with t(3;8)(p21;q12) translocations. Nat Genet 15(2):170–174.

12. Juma AR, Damdimopoulou PE, Grommen SV, Van de Ven WJ, & De Groef B (2016) Emerging role of PLAG1 as a regulator of growth and reproduction. J Endocrinol 228(2):R45–56.

13. Tohonen V, et al. (2015) Novel PRD-like homeodomain transcription factors and retrotransposon elements in early human development. Nat Commun 6:8207.

14. Hensen K, et al. (2004) Targeted disruption of the murine Plag1 proto-oncogene causes growth retardation and reduced fertility. Dev Growth Differ 46(5):459–470.

15. Ullu E & Tschudi C (1984) Alu sequences are processed 7SL RNA genes. Nature 312(5990):171–172.

16. Labuda D, Sinnett D, Richer C, Deragon JM, & Striker G (1991) Evolution of mouse B1 repeats: 7SL RNA folding pattern conserved. J Mol Evol 32(5):405–414.

17. Voz ML, Agten NS, Van de Ven WJ, & Kas K (2000) PLAG1, the main translocation target in pleomorphic adenoma of the salivary glands, is a positive regulator of IGF-II. Cancer Res 60(1):106–113.

18. Fantl V, Stamp G, Andrews A, Rosewell I, & Dickson C (1995) Mice lacking cyclin D1 are small and show defects in eye and mammary gland development. Genes Dev 9(19):2364–2372.

19. Lefebvre L, et al. (1998) Abnormal maternal behaviour and growth retardation associated with loss of the imprinted gene Mest. Nat Genet 20(2):163–169.

20. Argente MJ, Calle EW, Garcia ML, & Blasco A (2017) Correlated response in litter size components in rabbits selected for litter size variability. J Anim Breed Genet 134(6):505–511.

21. Bruce NW & Wellstead JR (1992) Spacing of fetuses and local competition in strains of mice with large, medium and small litters. J Reprod Fertil 95(3):783–789.

22. Vogt EJ, Meglicki M, Hartung KI, Borsuk E, & Behr R (2012) Importance of the pluripotency factor LIN28 in the mammalian nucleolus during early embryonic development. Development 139(24):4514–4523.

23. Tong ZB, et al. (2000) Mater, a maternal effect gene required for early embryonic development in mice. Nat Genet 26(3):267–268.

24. Van Dyck F, et al. (2007) Loss of the PlagL2 transcription factor affects lacteal uptake of chylomicrons. Cell Metab 6(5):406–413.

25. Wong CC, et al. (2010) Non-invasive imaging of human embryos before embryonic genome activation predicts development to the blastocyst stage. Nat Biotechnol 28(10):1115–1121.

26. Wang H & Dey SK (2006) Roadmap to embryo implantation: clues from mouse models. Nat Rev Genet 7(3): 185–199.

27. Elbarbary RA, Lucas BA, & Maquat LE (2016) Retrotransposons as regulators of gene expression. Science 351(6274):aac7247.

28. Mouse Genome Sequencing C, et al. (2002) Initial sequencing and comparative analysis of the mouse genome. Nature 420(6915):520–562.

29. Polak P & Domany E (2006) Alu elements contain many binding sites for transcription factors and may play a role in regulation of developmental processes. BMC Genomics 7:133.

30. Ge SX (2017) Exploratory bioinformatics investigation reveals importance of “junk” DNA in early embryo development. BMC Genomics 18(1):200.

31. Pelletier J, Thomas G, & Volarevic S (2018) Ribosome biogenesis in cancer: new players and therapeutic avenues. Nat Rev Cancer 18(1):51–63.

32. Fink T, et al. (2017) Functional confirmation of PLAG1 as the candidate causative gene underlying major pleiotropic effects on body weight and milk characteristics. Sci Rep 7:44793.

33. Rubin CJ, et al. (2012) Strong signatures of selection in the domestic pig genome. Proc Natl Acad Sci U S A 109(48):19529–19536.

34. Utsunomiya YT, et al. (2013) Genome-wide association study for birth weight in Nellore cattle points to previously described orthologous genes affecting human and bovine height. BMC Genet 14:52.

35. Zhang W, et al. (2016) Multi-strategy genome-wide association studies identify the DCAF16-NCAPG region as a susceptibility locus for average daily gain in cattle. Sci Rep 6:38073.

36. Juma AR, et al. (2017) PLAG1 deficiency impairs spermatogenesis and sperm motility in mice. Sci Rep 7(1):5317.

37. Krjutskov K, et al. (2016) Single-cell transcriptome analysis of endometrial tissue. Hum Reprod 31(4):844–853.

38. Kim D, et al. (2013) TopHat2: accurate alignment of transcriptomes in the presence of insertions, deletions and gene fusions. Genome Biol 14(4):R36.

39. Heinz S, et al. (2010) Simple combinations of lineage-determining transcription factors prime cis-regulatory elements required for macrophage and B cell identities. Mol Cell 38(4):576–589.

40. R Development Core Team (2010) R: A language and environment for statistical computing (R Foundation for Statistical Computing, Vienna, Austria).

41. Robinson MD, McCarthy DJ, & Smyth GK (2010) edgeR: a Bioconductor package for differential expression analysis of digital gene expression data. Bioinformatics 26(1):139–140.

42. Gu Z, Eils R, & Schlesner M (2016) Complex heatmaps reveal patterns and correlations in multidimensional genomic data. Bioinformatics 32(18):2847–2849.

43. Trapnell C, et al. (2014) The dynamics and regulators of cell fate decisions are revealed by pseudotemporal ordering of single cells. Nat Biotechnol 32(4):381–386.

44. Yu G, et al. (2010) GOSemSim: an R package for measuring semantic similarity among GO terms and gene products. Bioinformatics 26(7):976–978.

45. Ritchie ME, et al. (2015) limma powers differential expression analyses for RNA-sequencing and microarray studies. Nucleic Acids Res 43(7):e47.

46. Gupta S, Stamatoyannopoulos JA, Bailey TL, & Noble WS (2007) Quantifying similarity between motifs. Genome Biol 8(2):R24.

47. Bailey TL & Elkan C (1994) Fitting a mixture model by expectation maximization to discover motifs in biopolymers. Proc Int Conf Intell Syst Mol Biol 2:28–36.

48. Meng X, Brodsky MH, & Wolfe SA (2005) A bacterial one-hybrid system for determining the DNA-binding specificity of transcription factors. Nat Biotechnol 23(8):988–994.

49. Sandelin A, Alkema W, Engstrom P, Wasserman WW, & Lenhard B (2004) JASPAR: an open-access database for eukaryotic transcription factor binding profiles. Nucleic Acids Res 32(Database issue):D91–94.

50. Bailey TL & Gribskov M (1998) Combining evidence using p-values: application to sequence homology searches. Bioinformatics 14(1):48–54.

51. Hubley R, et al. (2016) The Dfam database of repetitive DNA families. Nucleic Acids Res 44(D1):D81–89.

