## Supplementary material for "Pleomorphic Adenoma Gene 1 Is Needed For Timely Zygotic Genome Activation and Early Embryo Development"

**INDEX**

**Video S1 (a separate attachment), legend** **Page 3**

Time-lapse video of WT embryo development

**Video S2 (a separate attachment), legend** **Page 3**

Time-lapse video of *matPlag1KO* embryo development

**File S1 (a separate attachment), legend** **Page 3**
*de novo* PLAG1 motifs and repetitive elements in human ZGA genes

**File S2 (a separate attachment), legend** **Page 3**
GO clusters associated with delayed-activation and delayed-degradation genes

**File S3 (a separate attachment), legend** **Page 4**
GO clusters associated with up- and downregulated genes during ZGA in WT mice

**File S4 (a separate attachment), legend** **Page 4**
*de novo* PLAG1 motifs in mouse ZGA genes

**Figure S1** **Page 4**
Comparison of human PLAG1 and mouse Plag1 protein sequence

**Figure S2** **Page 5**

*Plag1* deficiency does not have significant effects on ovaries and uterus

**Figure S3** **Page 6**
Size of the mRNA and spike-in reference RNA libraries

**Figure S4** **Page 7**

Expression of *PLAG1* in human embryos

**Figure S5** **Page 8**
Expression of *Plag1* transcripts in mouse ovary as shown by X-gal staining

**Figure S6** **Page 9**

Gene set comparison of mouse and human ZGA genes

**Figure S7** **Page 10**

Frequency of *de novo* PLAG1 motifs and B1 elements in delayed-activation and delayed-degradation gene promoters

**References** **Page 11**

**Video S1.** Time-lapse video of wildtype (WT) embryo development

A representative movie of WT mouse embryo development from zygote to morula. The video time line is indicated (0-89 h) as well as the time spent in 2-cell stage.

**Video S2.** Time-lapse video of *matPlag1KO* embryo development

A representative movie of *matPlag1KO* mouse embryo development from zygote to morula. The video time line is indicated (0-89 h) as well as the time spent in 2-cell stage.

**File S1.** *de novo* PLAG1 motifs and repetitive elements in human zygotic genome activation (ZGA) genes

<u>Sheet MEME</u> Position of source sequences for the *de novo* PLAG1 motif in human ZGA transcript far 5' ends (TFEs) within –2,000 bp upstream to 500 bp downstream from the transcription start site (TSS). The sheet gives the MEME (Bailey and Elkan 1994) analysis results for human ZGA TFEs that were used to derive the *de novo* motif that is similar to the PLAG1 binding motif MA0163.1 (Meng et al. 2005) in JASPAR (Sandelin et al. 2004). Source sequence position, source strand, p-value, TFE region, promoter region, TFE and promoter strand, associated gene and TFE position within the gene are shown, as well as the mouse homologs and their developmental function as annotated in the database of transcriptome in mouse early embryos (DBTMEE). The concept TFE, an identifier of transcription start site, was used in our earlier human ZGA study (Tohonen et al. 2015).

<u>Sheet MAST</u> Position of sequences similar to the *de novo* PLAG1 motif in the human ZGA TFEs within –2,000 bp upstream to 500 bp downstream from their transcription start site (TSS). The sheet gives the MAST (Bailey and Gribskov 1998) analysis results for the 93 TFEs that contain sites similar to the *de novo* PLAG1 motif. The position of the motif, motif strand, p-value, TFE region, promoter region, TFE and promoter strand, associated gene, and TFE position within the gene are shown.

<u>Sheet AluJSY</u> Position of Alu elements in the human ZGA TFEs within –2,000 bp upstream to 500 bp downstream from their transcription start site (TSS). AluS/J/Y elements were extracted from the RepeatMasker track of the UCSC Genome Browser, and the columns about TFE were joined. The type of element, its position ("left" and "right" within the promoter 0-2,500), TFE region, promoter region, TFE and promoter strand, associated gene, and TFE position within the gene are shown.

**File S2.** Gene ontology (GO) clusters associated with delayed-activation and delayed-degradation genes

Genes affected by maternal *Plag1* deficiency in 2-cell mouse embryos were assigned to GO categories using topGO library in R (R Development Core Team 2010) using the classic algorithm and Fisher statistics. Top 150 GOs by p-value were then clustered based on their semantic similarity. Clusters 1-8 (delayed up) and 1-6 (delayed down) are shown.

**File S3.** Gene ontology (GO) clusters associated with up- and downregulated genes during zygotic genome activation (ZGA) in wildtype (WT) mice

Genes up- and downregulated during major ZGA in wild type mouse embryos (2-cell to 8-cell transition) were assigned to GO categories using topGO library in R (R Development Core Team 2010) using the classic algorithm and Fisher statistics. Top 150 GOs by p-value were then clustered based on their semantic similarity. Clusters 1-9 are shown.

**File S4.** *de novo* PLAG1 motifs in mouse zygotic genome activation (ZGA) genes

All annotated mouse promoters were scanned for the presence of the *de novo* PLAG1 motif from –2,000 bp upstream to 500 bp downstream of transcriptional start sites (TSSs) using Homer (Heinz et al. 2010). Delayed-activation genes that have *de novo* PLAG1 motifs in their promoters are shown (symbol, entrez, refSeq) together with the motif sequence, strand and genomic coordinates of the motif as well as the location of the transcription start site (TSS) and the distance relative to the direction of the gene. The last column indicates if the gene is also present among the human ZGA genes (Tohonen et al. 2015).

**Figure S1.** Comparison of human PLAG1 and mouse Plag1 protein sequence

```
                                             +-------------------1-+
                     1         2         3         4         5         6
            123456789012345678901234567890123456789012345678901234567890
Human  1    MATVIPGDLSEVRDTQKVPSGKRKRGETKPRKNFPCQLCDKAFNSVEKLKVHSYSHTGER  60
            MATVIPGDLSEVRDTQK PSGKRKRGE+KPRKNFPCQLCDKAFNSVEKLKVHS+SHTGER
Mouse  1    MATVIPGDLSEVRDTQKAPSGKRKRGESKPRKNFPCQLCDKAFNSVEKLKVHSFSHTGER  60

              +---------------------2-+    +-------------------3-+
            6        7         8         9        10        11        12
            123456789012345678901234567890123456789012345678901234567890
Human  61   PYKCIQQDCTKAFVSKYKLQRHMATHSPEKTHKCNYCEKMFHRKDHLKNHLHTHDPNKET  120
            PYKC  QDCTKAFVSKYKLQRHMATHSPEKTHKCNYCEKMFHRKDHLKNHLHTHDPNKET
Mouse  61   PYKCTHQDCTKAFVSKYKLQRHMATHSPEKTHKCNYCEKMFHRKDHLKNHLHTHDPNKET  120

            +--------------------4-+     +--------------------5-+
           12        13        14        15        16        17        18
            123456789012345678901234567890123456789012345678901234567890
Human  121  FKCEECGKNYNTKLGFKRHLALHAATSGDLTCKVCLQTFESTGVLLEHLKSHAGKSSGGV  180
            FKCEECGK+YNTKLGFKRHLALHAATSGDLTCKVCLQ FESTGVLLEHLKSHAGKSSGGV
Mouse  121  FKCEECGKSYNTKLGFKRHLALHAATSGDLTCKVCLQNFESTGVLLEHLKSHAGKSSGGV  180

                +--------------------6-+    +---------------------7-+
           18        19        20        21        22        23        24
            123456789012345678901234567890123456789012345678901234567890
Human  181  KEKKHQCEHCDRRFYTRKDVRRHMVVHTGRKDFLCQYCAQRFGRKDHLTRHMKKSHNQEL  240
            KEKKHQCEHC+RRFYTRKDVRRHMVVHTGRKDFLCQYCAQRFGRKDHLTRHMKKSHNQEL
Mouse  181  KEKKHQCEHCERRFYTRKDVRRHMVVHTGRKDFLCQYCAQRFGRKDHLTRHMKKSHNQEL  240

            123456789012345678901234567890123456789012345678901234567890
Human  241  LKVKTEPVDFLDPFTCNVSVPIKDELLPVMSLPSSELLSKPFTNTLQLNLYNTPFQSMQS  300
            LKVKTEPVDFLDPFTCN+SVPIKDELLPVMSLPSSELLSKPFTNTLQLNLYNTPFQSMQS
Mouse  241  LKVKTEPVDFLDPFTCNMSVPIKDELLPVMSLPSSELLSKPFTNTLQLNLYNTPFQSMQS  300

Human  301  SGSAHQMITTLPLGMTCPIDMDTVHPSHHLSFKYPFSSTSYAISIPEKEQPLKGEIESYL  360
            SGSAHQMITTLPLGMTCPIDMD VHPSHHL+FK PFSSTSYAISIPEKEQPLKGEIESYL
Mouse  301  SGSAHQMITTLPLGMTCPIDMDAVHPSHHLAFKCPFSSTSYAISIPEKEQPLKGEIESYL  360

Human  361  MELQGGVPsssqdsqassssKLGLDPQIGSLDDGAGDlslskssisisDPLNTPALDFSQ  420
            MELQGG P SS     +SSSKLGL+PQ GS DDGAGDLSLSKSSISISDPL+TPALDFSQ
Mouse  361  MELQGGAP-SSSQDSPASSSKLGLEPQSGSPDDGAGDLSLSKSSISISDPLSTPALDFSQ  419

Human  421  LFNFIPLNGPPYNPLSVGSLGMSYSQEEAHSSVSqlppqtqdlqdpANTIGLGslhslsa  480
            LFNFIPLNGPPYNPLSVGSLGMSYSQEEAHSSVSQLP QTQDLQDPANT+GL SLHSLSA
Mouse  420  LFNFIPLNGPPYNPLSVGSLGMSYSQEEAHSSVSQLPTQTQDLQDPANTVGLSSLHSLSA  479

Human  481  aftsslststtlPRFHQAFQ  500
            AFTSSLS+STTLPRFHQAFQ
Mouse  480  AFTSSLSSSTTLPRFHQAFQ  499
```

Comparison of amino acid sequences between human PLAG1 (RefSeq NP_002646.2) and mouse PLAG1 (NP_064353.2) by NCBI blastp (Altschul et al. 1990) indicating 94% similarity. The seven C2H2 zinc-finger domains are labeled (+---+) based on UniProt protein knowledge base (Pundir et al. 2017) (entry Q6DJT9). Domains 6 and 7 bind to the “core” of the PLAG1 consensus sequence while domain 3 binds to the “cluster” [Figure 1c *(v)* and (Voz et al. 2000)]. The amino acid sequences of these three domains are identical between mice and humans except for position 191 (red) within domain 6. However, human PLAGL1 has a glutamic acid residue (E) at the corresponding position of the PLAG1 D191E, and binding preference to G-rich “core” was highly conserved in both of PLAG1 and PLAGL1 (Hensen et al. 2002). Moreover, similarity of the C2H2 domains of mouse PLAG1 to human PLAG1 is higher than human PLAGL1. Therefore, although mouse PLAG1 has one inconsistent residue within the C2H2 domain for binding, preference of the binding site sequences would be identical between human and mouse PLAG1.

**Figure S2.** *Plag1* deficiency does not have significant effects on ovaries and uterus.

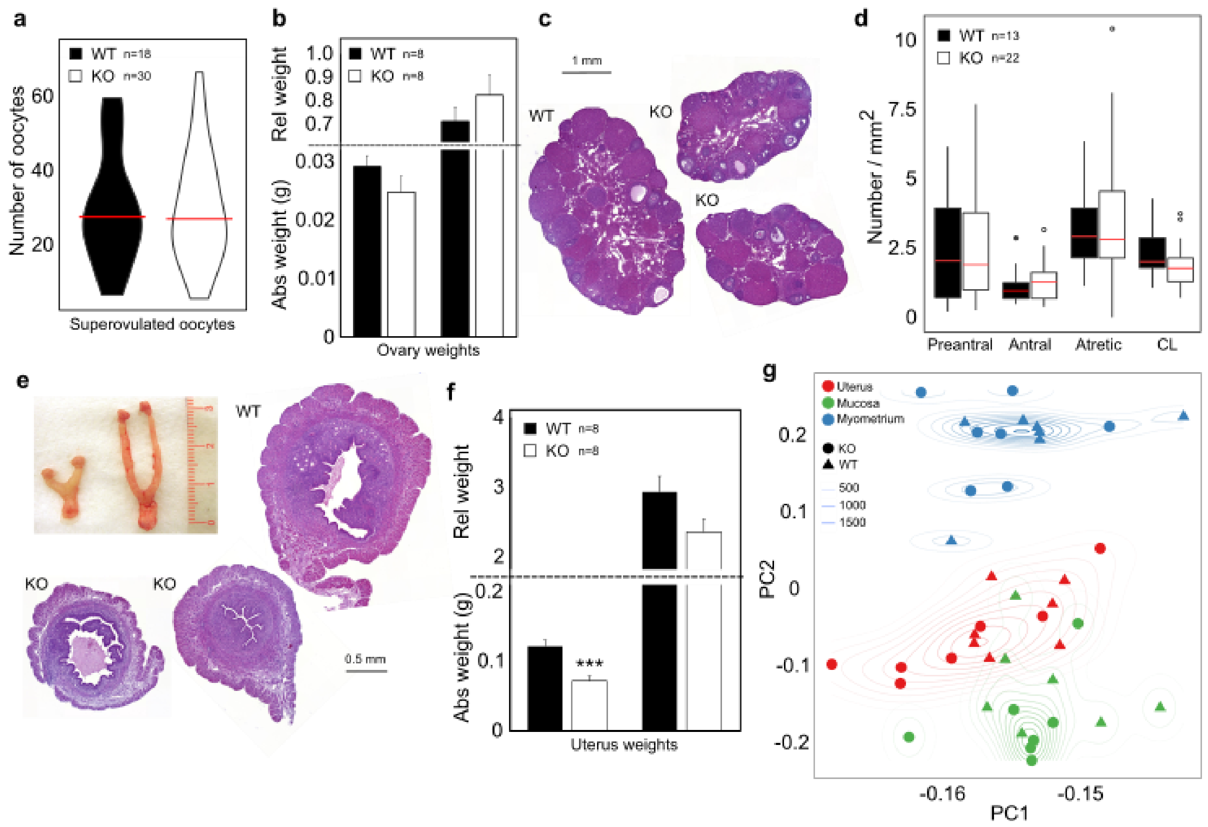


**(a)** Number of oocytes collected after superovulation presented as violin plots. The median is marked as a red line. **(b)** Absolute and relative (body weight-adjusted) weights of ovaries, shown as means + SEM. **(c)** Representative images of hematoxylin-eosin-stained ovarian cross sections from wildtype (WT) and *Plag1* knockout (KO) mice. The scale bar is 1 mm. **(d)** Numbers of preantral, antral and atretic follicles as well as corpora lutea (CL) per mm$^2$ of ovary. Data are presented as median (red line), interquartile ranges (box), and non-outlier ranges (whisker). Outliers are depicted as dots. **(e)** Representative photos and hematoxylin-eosin-stained histological images of WT and *Plag1* KO uteri. The scale bar is 500 µm. **(f)** Absolute and relative (body weight-adjusted) uterine weights presented as means + SEM. **(g)** Principal component (PC) analysis of global gene expression in WT and KO uterus samples (N = 7) divided into “uterus” (a piece of uterine horn), “mucosa” (cells scraped from the endometrial surface of the uterine horn), and “myometrium” (tissue left after mucosa has been removed).
Statistical analysis by *t*-test (a, b, f) or two-way ANOVA (d); ***p < 0.001.

**Figure S3.** Size of the mRNA and spike-in reference RNA libraries

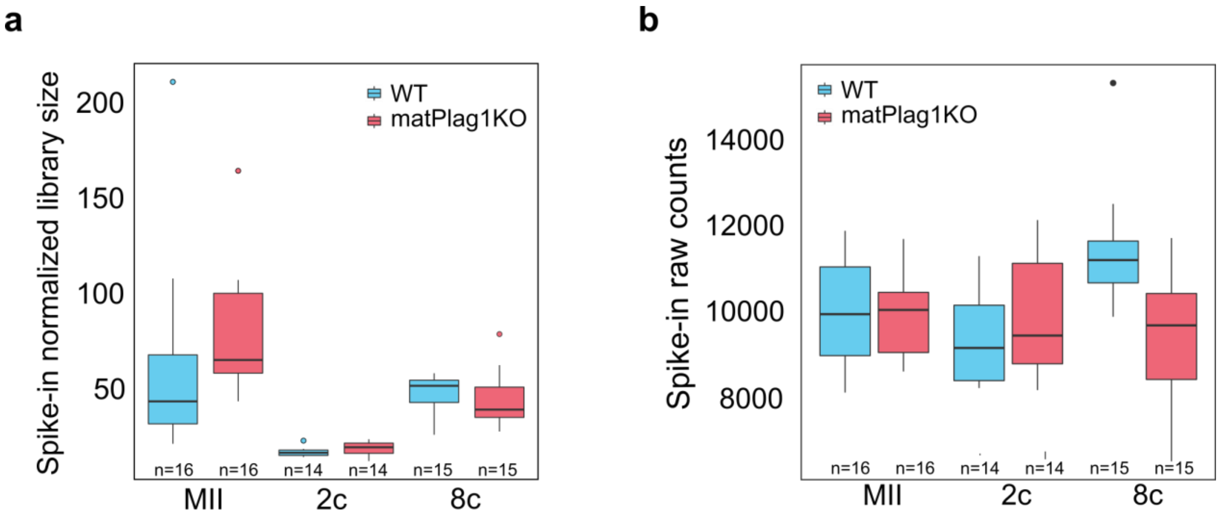


The size of the **(a)** spike-in normalized mRNA libraries and **(b)** spike-in reference RNA libraries recovered through RNA-sequencing. The size did not significantly vary between genotypes. MII, MII oocyte; 2c, 2-cell stage embryo; 8c, 8-cell stage embryo.

**Figure S4.** Expression of *PLAG1* in human embryos

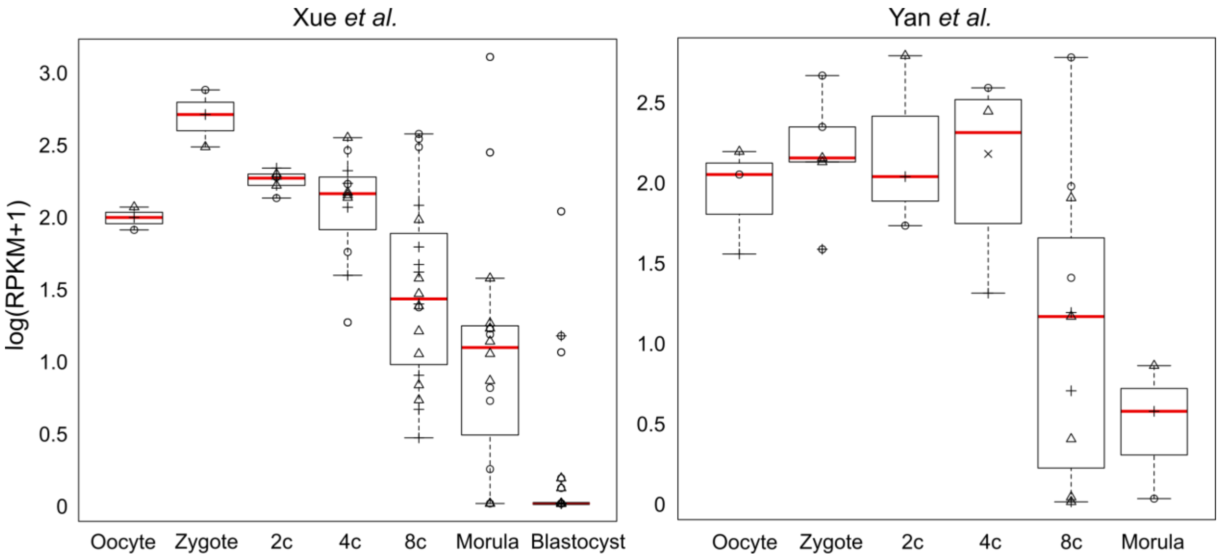


*PLAG1* expression in human pre-implantation embryos in two independent single-cell RNA-sequencing datasets. The normalized expression matrices were downloaded from GEO with the accession numbers GSE44183 (Xue et al. 2013) and GSE36552 (Yan et al. 2013). Expression is shown as log-transformed RPKM values by developmental stage. A constant (1) was added to the RPKM values prior to log-transformation to avoid negative values. The data are presented as box plots where median is shown as a red line, edges of the box are the upper and lower quartiles, and the whiskers show the highest and lowest values while excluding outliers. Cells from the same embryo in each developmental stage are depicted with the same shape. 2c, 2-cell; 4c, 4-cell; 8c, 8-cell; RPKM, reads per kilobase million.

**Figure S5**. Expression of *Plag1* transcripts in mouse ovary as shown by X-gal staining

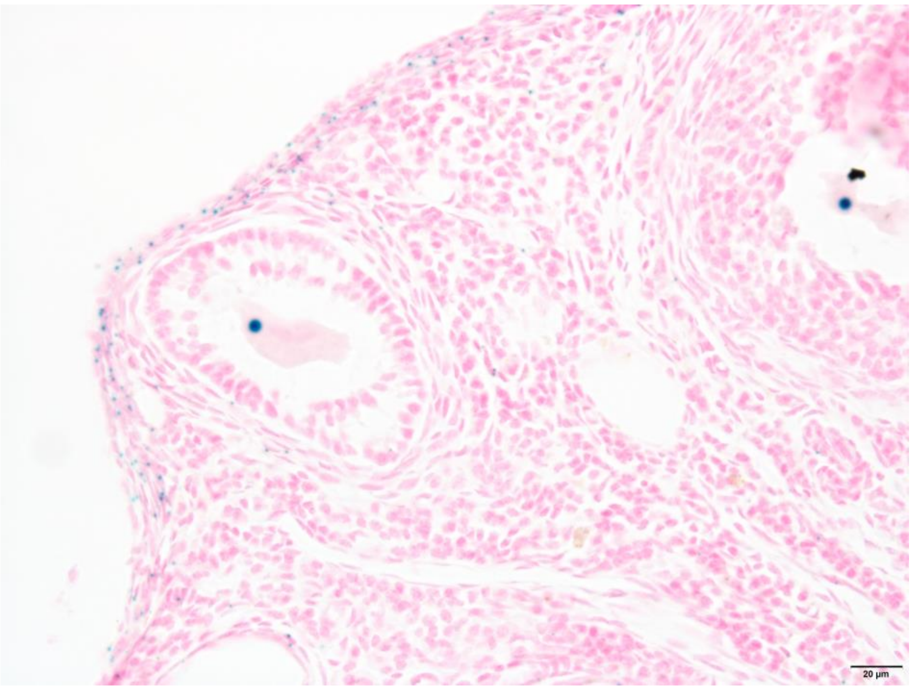


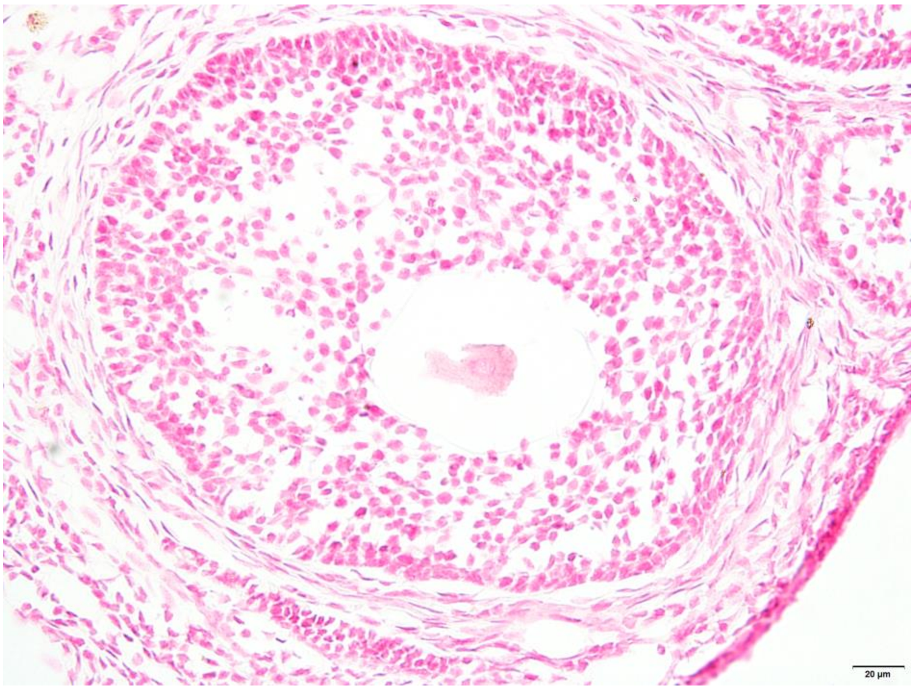


The mutant allele in the *Plag1KO* animals consists of the *lacZ* gene that replaces the entire open reading frame of *Plag1*, enabling visualization of *Plag1* promoter activity using X-gal staining. The top picture displays *lacZ*-positive cells (blue) in a representative ovary section of a *Plag1KO* mouse, aged between 4-5 months, as determined by X-gal staining. Expression of *lacZ* was found mostly in the nuclei of growing oocytes and in the tunica albugina. The bottom picture shows a negative control for X-gal staining (WT mouse), showing no blue signal. The scale bars are 20 μm.

**Figure S6**. Gene set comparison of mouse and human zygotic genome activation (ZGA) genes

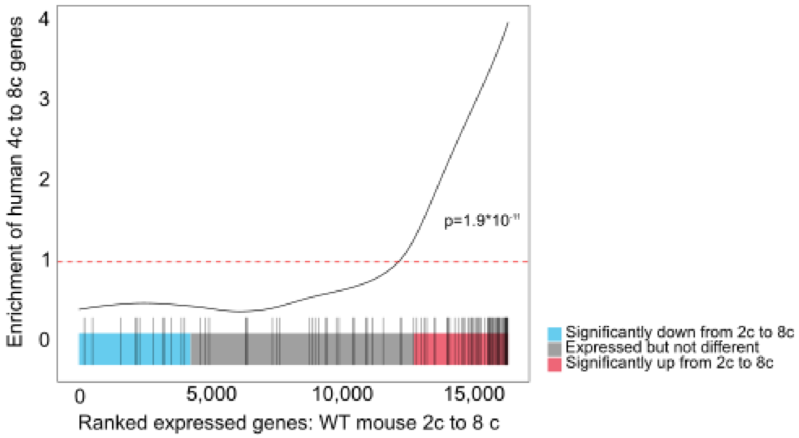


Gene set enrichment analysis comparing gene expression changes during mouse and human ZGA. Mouse ZGA genes are displayed on the x-axis ranked by the level of expression change, and human ZGA gene that have a mouse orthologue are depicted with black vertical lines. Enrichment of the human genes among the mouse genes is depicted wth the black line. No enrichment level (=1) is shown as red dotted line. Significance of the enrichment was tested with the GeneSet test function.

**Figure S7**. Frequency of *de novo* PLAG1 motifs and B1 elements in delayed-activation and delayed-degradation gene promoters.

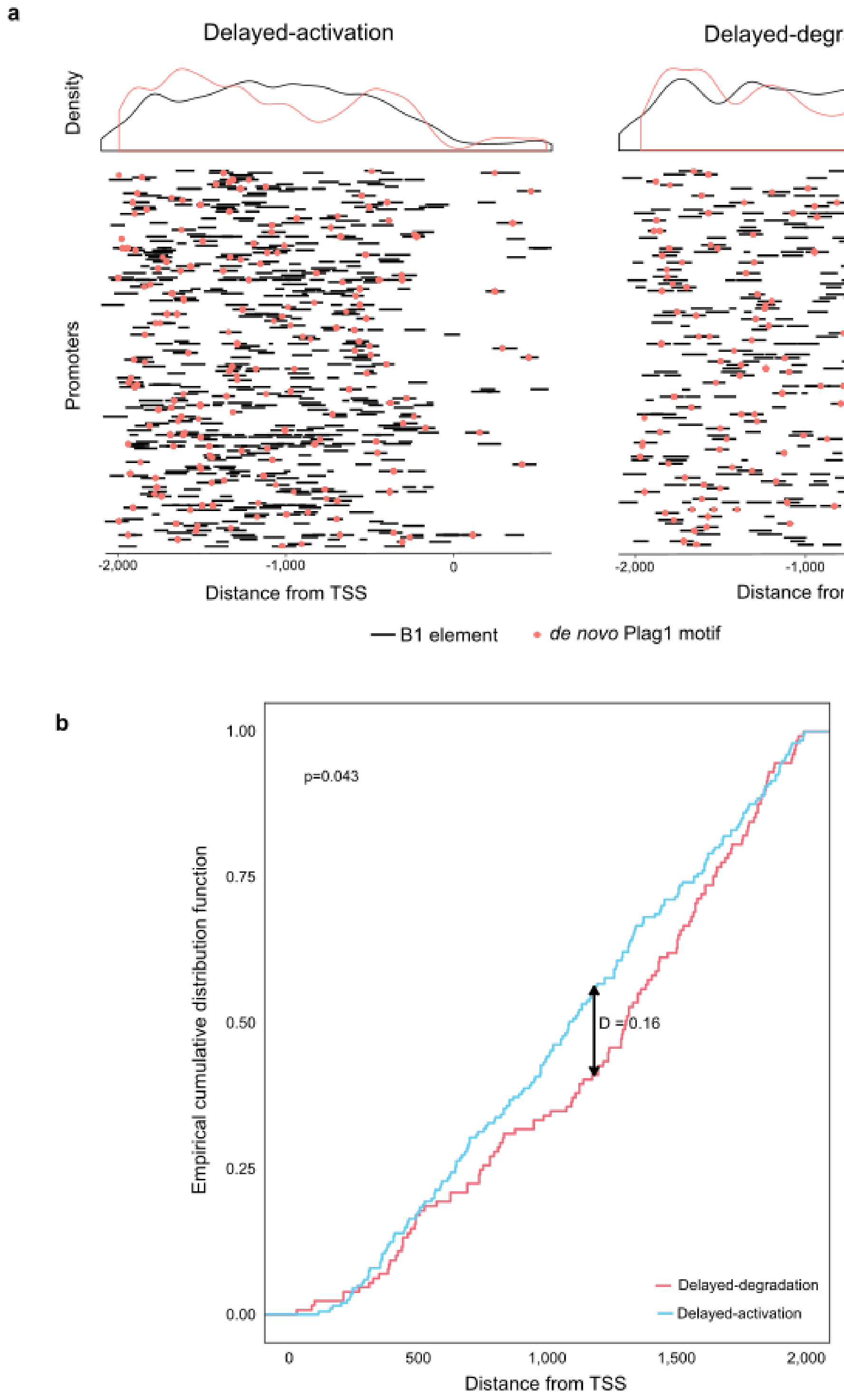


**(a)** Location of B1 repetitive elements (black lines) and *de novo* PLAG1 motifs (red dots) along the promoters of delayed-activation and delayed-degradation genes from -2,000 bp to +500 bp around the transcription start site (TSS). Enrichment of the sites as shown as color-coded lines above the graphs.
**(b)** Empirical cumulative distribution function showing the distribution of *de novo* PLAG1 motifs along the promoters of delayed-activation and delayed-degradation genes.

### References

Altschul SF, Gish W, Miller W, Myers EW, Lipman DJ. 1990. Basic local alignment search tool. *J Mol Biol* **215**: 403-410.

Bailey TL, Elkan C. 1994. Fitting a mixture model by expectation maximization to discover motifs in biopolymers. *Proc Int Conf Intell Syst Mol Biol* **2**: 28-36.

Bailey TL, Gribskov M. 1998. Combining evidence using p-values: application to sequence homology searches. *Bioinformatics* **14**: 48-54.

Heinz S, Benner C, Spann N, Bertolino E, Lin YC, Laslo P, Cheng JX, Murre C, Singh H, Glass CK. 2010. Simple combinations of lineage-determining transcription factors prime cis-regulatory elements required for macrophage and B cell identities. *Mol Cell* **38**: 576-589.

Hensen K, Van Valckenborgh IC, Kas K, Van de Ven WJ, Voz ML. 2002. The tumorigenic diversity of the three PLAG family members is associated with different DNA binding capacities. *Cancer Res* **62**: 1510-1517.

Meng X, Brodsky MH, Wolfe SA. 2005. A bacterial one-hybrid system for determining the DNA-binding specificity of transcription factors. *Nat Biotechnol* **23**: 988-994.

Pundir S, Martin MJ, O'Donovan C. 2017. UniProt Protein Knowledgebase. *Methods Mol Biol* **1558**: 41-55.

R Development Core Team. 2010. R: A language and environment for statistical computing. R Foundation for Statistical Computing, Vienna, Austria.

Sandelin A, Alkema W, Engstrom P, Wasserman WW, Lenhard B. 2004. JASPAR: an open-access database for eukaryotic transcription factor binding profiles. *Nucleic Acids Res* **32**: D91-94.

Tohonen V, Katayama S, Vesterlund L, Jouhilahti EM, Sheikhi M, Madissoon E, Filippini-Cattaneo G, Jaconi M, Johnsson A, Burglin TR et al. 2015. Novel PRD-like homeodomain transcription factors and retrotransposon elements in early human development. *Nat Commun* **6**: 8207.

Voz ML, Agten NS, Van de Ven WJ, Kas K. 2000. PLAG1, the main translocation target in pleomorphic adenoma of the salivary glands, is a positive regulator of IGF-II. *Cancer Res* **60**: 106-113.

Xue Z, Huang K, Cai C, Cai L, Jiang CY, Feng Y, Liu Z, Zeng Q, Cheng L, Sun YE et al. 2013. Genetic programs in human and mouse early embryos revealed by single-cell RNA sequencing. *Nature* **500**: 593-597.

Yan L, Yang M, Guo H, Yang L, Wu J, Li R, Liu P, Lian Y, Zheng X, Yan J et al. 2013. Single-cell RNA-Seq profiling of human preimplantation embryos and embryonic stem cells. *Nat Struct Mol Biol* **20**: 1131-1139.
